# Structural Elucidation of a Protective B cell Epitope on Outer Surface Protein C (OspC) of the Lyme disease spirochete, *Borreliella burgdorferi*

**DOI:** 10.1101/2022.11.28.518297

**Authors:** Michael J. Rudolph, Simon A. Davis, H M Emranul Haque, David D. Weis, David J Vance, Carol Lyn Piazza, Monir Ejemel, Lisa Cavacini, Yang Wang, M. Lamine Mbow, Robert D. Gilmore, Nicholas J Mantis

## Abstract

<u>O</u>uter <u>s</u>urface <u>p</u>rotein C (OspC) plays a pivotal role in mediating tick-to-host transmission and infectivity of the Lyme disease spirochete, *Borreliella burgdorferi*. OspC is a helical-rich homodimer that interacts with tick salivary proteins, as well as components of the mammalian immune system. Several decades ago, it was shown that the OspC-specific monoclonal antibody, B5, was able to passively protect mice from experimental tick-transmitted infection by *B. burgdorferi* strain B31. However, B5’s epitope has never been elucidated, despite widespread interest in OspC as a possible Lyme disease vaccine antigen. Here we report the crystal structure of B5 antigen-binding fragments (Fabs) in complex with recombinant OspC type A (OspC_A_). Each OspC monomer within the homodimer was bound by a single B5 Fab in a side-on orientation, with contact points along OspC’s α-helix 1 and α-helix 6, as well as interactions with the loop between a-helices 5 and 6. In addition, B5’s complementarity-determining region (CDR) H3 bridged the OspC-OspC’ homodimer interface, revealing the quaternary nature of the protective epitope. To provide insight into the molecular basis of B5 serotype specificity, we solved the crystal structures of recombinant OspC types B and K and compared them to OspC_A_. This study represents the first structure of a protective B cell epitope on OspC and will aid in the rational design of OspC-based vaccines and therapeutics for Lyme disease.

**IMPORTANCE:** The spirochete, *Borreliella burgdorferi*, is the causative agent of Lyme borreliosis, the most common tickborne disease in the United States. The spirochete is transmitted to humans during the course of a tick taking a bloodmeal. After *B. burgdorferi* is deposited into the skin of a human host, it replicates locally and spreads systemically, often resulting in clinical manifestations involving the central nervous system, joints, and/or heart. Antibodies directed against *B. burgdorferi*’s outer surface protein C (OspC) are known to block tick-to-host transmission, as well as dissemination of the spirochete within a mammalian host. In this report, we reveal the first atomic structure of one such antibody in complex with OspC. Our results have implications for the design of a Lyme disease vaccine capable to interfering with multiple stages in *B. burgdorferi* infection.

## Introduction

*Borreliella burgdorferi* is the principal etiological agent of Lyme disease (LD), the most common tick-borne infection in North America and Europe. In the United States, the vector of *B. burgdorferi* is the blacklegged ticks, *Ixodes scapularis* and *Ixodes pacificus*. Naïve larvae acquire *B. burgdorferi* during a blood meal on an infected reservoir species, including birds and an array of small mammals. Once within a tick, the spirochetal bacteria remain dormant in the midgut until a second blood meal taken during the nymphal stage, after which *B. burgdorferi* migrates to the salivary glands where it is deposited into the skin of an impending host. Perpetuation of *B. burgdorferi*’s enzoonotic cycle and persistence within vertebrate reservoirs is contingent on the spirochete’s ability to evade innate and adaptive immunity (Di et al., 2022; Rana et al., 2022). Although humans are incidental (“dead end”) hosts for *B. burgdorferi*, the spirochetes have the capacity to disseminate and colonize distal tissues, which contributes to the clinical manifestations commonly associated with Lyme disease, including erythema migrans, neuroborreliosis, carditis, and Lyme arthritis (Radolf et al., 2021; Steere et al., 2016). With an estimated 476,000 cases of Lyme disease per year in the United States (Kugeler et al., 2021), there is an urgent need for vaccines that prevent *B. burgdorferi* transmission and limit human infections (Dattwyler and Gomes-Solecki, 2022; Gomes-Solecki et al., 2020; Wormser, 2022).

Outer surface protein C (OspC) has long been considered a candidate Lyme disease vaccine antigen. OspC is a ~23 kDa helical-rich lipoprotein expressed by *B. burgdorferi* in the tick midgut during the course of a blood meal (Caimano et al., 2019; Eicken et al., 2001; Kumaran et al., 2001; Ohnishi et al., 2001; Schwan et al., 1995; Zuckert et al., 2001). OspC facilitates migration of *B. burgdorferi* from the midgut to the salivary glands and contributes to early stages of mammalian survival (De Silva and Fikrig, 1995; Ohnishi *et al*., 2001; Pal et al., 2004; Schwan *et al*., 1995). In the mouse model, OspC antibodies elicited through active or passive vaccination prevent tick-mediated *B. burgdorferi* infection (Gilmore et al., 1996; Mbow et al., 1999). Recent evidence indicates OspC antibodies entrap *B. burgdorferi* within the tick midgut by a mechanism that does not involve direct spirochete killing (Bhatia et al., 2018). OspC antibodies also protect mice against *B. burgdorferi* needle infection and have been reported to promote resolution of arthritis and carditis in SCID mouse model of Lyme disease (Gilmore *et al*., 1996; Preac-Mursic et al., 1992; Probert and LeFebvre, 1994; Zhong et al., 1997). Thus,

OspC antibodies afford a double layer of protection by interfering with *B. burgdorferi* tick-to-host transmission and promoting clearance of spirochetes that do gain access to a mammalian host.

A major limitation of OspC as a vaccine antigen is its polymorphism within and across *B. burgdorferi* genospecies. At least 25 different *ospC* alleles or types have been identified to date, with extensive diversity occurring even within limited geographical regions (Baum et al., 2013; Bunikis et al., 2004; Earnhart et al., 2005; Hanincova et al., 2013; Seinost et al., 1999; Wang et al., 1999). The degree of OspC variability is such that antibody responses to a given OspC type have limited cross-reactivity with other OspC types (Barbour and Travinsky, 2010; Bhatia *et al*., 2018; Bockenstedt et al., 1997). This is consequential because the immunodomnant nature of OspC and that susceptibility to *B. burgdorferi* re-infection and repeated episodes of Lyme disease has been linked to OspC variability (Baum *et al*., 2013; Bhatia *et al*., 2018; Bockenstedt *et al*., 1997; Di *et al*., 2022; Gilmore *et al*., 1996; Nadelman et al., 2012; Rousselle et al., 1998). Thus, a vaccine based on a single OspC type would have limited utility in regions where multiple *B. burgdorferi* OspC types co-exist.

A multivalent or chimeric OspC vaccine consisting of conserved or minimally variable epitopes might afford protection against diverse *B. burgdorferi* strains (Izac and Marconi, 2019). The structures of OspC types A, I and E were solved two decades ago and revealed a high degree of tertiary and quaternary similarity (**Table 1; Figure S1**) (Eicken *et al*., 2001; Kumaran *et al*., 2001). In each case, OspC consisted of four long α-helices (1, 2, 3, 6), two shorter a-helices (4 and 5), and two short anti-parallel β-strands located between α1 and α2 (Eicken *et al*., 2001; Kumaran *et al*., 2001). The biologically functional molecule is a dimer (with monomers referred to as OspC-OspC’ herein) that assumes a knob-shaped structure anchored via N-terminal lipidated moieties in the spirochete’s outer surface membrane (Eicken *et al*., 2001; Kumaran *et al*., 2001; Zuckert *et al*., 2001). The C-terminus of OspC folds back onto the rest of the molecule such that α-helix 6 runs antiparallel to α-helix 1 and terminates at the bacterial surface. Multiple sequence alignment (MSA) of 23 OspC proteins revealed blocks of conserved residues localized in the N- (residues ~31-78) and C-termini (residues ~191-206), whereas high degrees of variably are within the short anti-parallel β-strands and other patches throughout the molecule (Baum *et al*., 2013; Earnhart *et al*., 2005; Earnhart et al., 2010; Earnhart and Marconi, 2007; Wilske et al., 1992). Computational modeling and linear B cell epitope analysis has identified regions on OspC likely to be the major determinants of type-specific and cross-reactive antibody recognition (Baum *et al*., 2013; Earnhart *et al*., 2005; Earnhart *et al*., 2010; Earnhart and Marconi, 2007; Izac et al., 2019; Izac et al., 2020; Wilske *et al*., 1992). However, no structure of a conformational (non-linear) protective epitope on OspC has been reported to date.

**Table 1.** OspC crystal structures and PDB codes.

| Table 1. OspC crystal structures and PDB codes |  |  |  |
| --- | --- | --- | --- |
| Type | Strain | PDB ID | Reference |
| A | B31 | 1GGQ | (Kumaran *et al.*, 2001) |
| B | ZS7 | 7UJ2 | this study |
| E | N40 | 1G5Z | (Eicken *et al.*, 2002) |
| I | HB19 | 1F1M | (Kumaran *et al.*, 2001) |
| K | 297 | 7UJ6 | this study |

B5 is an OspC-specific mouse monoclonal IgG antibody (MAb) first identified in a screen of B cell hybridomas from mice infected with *B. burgdorferi* strain B31 (Mbow et al., 2002). Passive immunization studies demonstrated that B5 IgG is sufficient to protect mice against experimental tick-mediated *B. burgdorferi* challenge, possibly by interfering with spirochete egress from the tick midgut (Gilmore and Piesman, 2000; Mbow *et al*., 1999). B5 IgG remains the only OspC MAb to have been shown to be protective against natural route of *B. burgdorferi* transmission, making it a valuable reagent to understand the molecular mechanism of OspC antibody function towards *B. burgdorferi*. In this study, we report the X-ray crystal of B5 Fab fragments in complex with an OspC_A_ homodimer. The structure represents the first elucidation of a protective epitope on OspC and unveils molecular interactions that constrain the breadth of B5 reactivity across OspC types.

## Results and Discussion

B5 IgG was originally isolated from mice that had been experimentally challenged with *B. burgdorferi* strain B31 (Gilmore and Mbow, 1999; Mbow *et al*., 1999). Due to limited supply of mouse B5 IgG, we generated a recombinant chimeric derivative of B5 in which murine V_H_ and V_L_ elements were fused to human IgG_1_ Fc and kappa light chains and expressed in Expi293 cells. Using purified mouse B5 IgG as well as the chimeric-derivative of B5 IgG_1_, we confirmed B5 reactivity with OspC. Recognition of native OspC on the surface of viable spirochetes was demonstrated by flow cytometry: a derivative *B. burgdorferi* strain B31 was incubated with B5 IgG or an isotype control (PB10 IgG) followed by Alexa647-labeled secondary antibody then analyzed for mean fluorescence intensity (MFI). B5 IgG labeled >80% of the live spirochetes with an MFI >3200, as compared to the isotype control, which labeled <1% of cells with MFI of ~50 (**Figure 1A,B**). To assess the breadth of B5 IgG reactivity, ELISA plates coated with recombinant OspC_A_, OspC type B (OspC_B_) and K (OspC_K_), then probed with a range of B5 IgG concentrations. By ELISA, B5 IgG reacted exclusively with OspC_A_, indicating that the MAb has limited OspC cross reactivity (**Figure 1C**). Similar results were observed by dot blot (**Figure 1D**). Using biolayer inferometry (BLI), B5 IgG had an apparent dissociation constant (*K*_D_) of ~ 0.2 nM for recombinant OspC_A_ (**Figure S2**). These results confirm that B5 IgG recognizes both native and recombinant OspC_A_.

**Figure 1.**
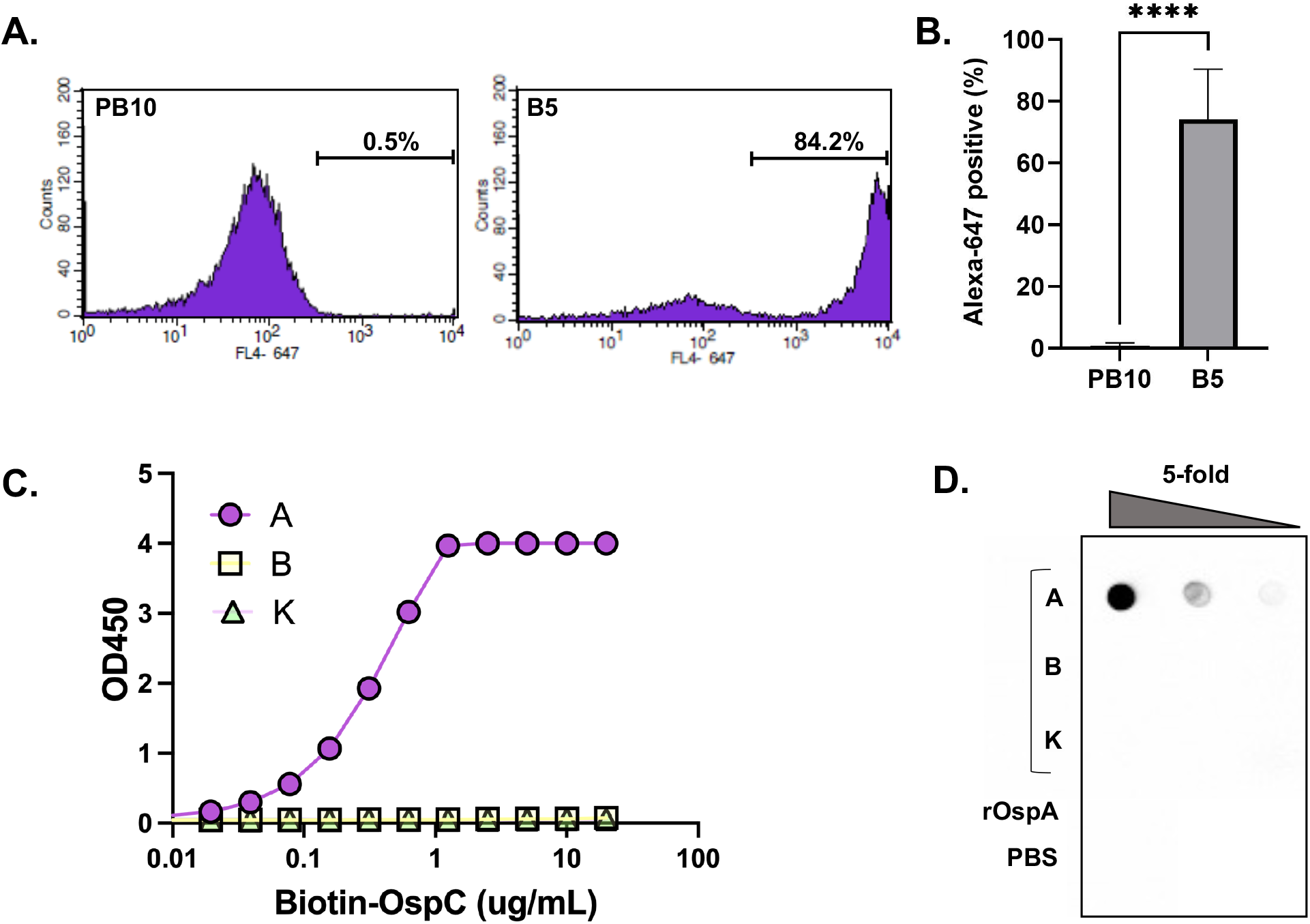
Reactivity of B5 IgG with native and recombinant OspC. (**A, B**). Mid-log phase *B. burgdorferi* over expressing OspC_A_ was incubated with chimeric B5 IgG or an isotype control (PB10) for 1 h at 37°C, washed and then labeled with Alexa 647-labeled goat anti-human secondary antibody. Cells were analyzed using a BD FACS Calibur flow cytometer. Panel A shows a representative histogram of PB10 (left) and B5 (right) with relative fluorescence plotted on x-axis and number of events plotted on y-axis. In panel B, the total number of Alexa-positive cells following PB10 and B5 treatment from three independent replicates. Asterisks indicate statistical differences (p <0.001) based on Student’s t-test. (**C**) Capture of biotin-tagged recombinant OspC types A, B and K by immobilized B5 IgG, as described in Material and Methods. The capture ELISA is representative of three independent replicates. (**D**) Reactivity of B5 IgG with 5-fold serial dilutions of recombinant OspC types A, B and K spotted onto nitrocellulose membrane. Recombinant OspA (rOspA) and PBS were spotted as negative controls.

### B5 IgG epitope localization using hydrogen exchange-mass spectrometry (HX-MS)

It was previously reported that B5 IgG recognizes a conformationally-sensitive epitope on OspC, possibly involving the C-terminus of α-helix 6 (Gilmore and Mbow, 1999). To localize B5 IgG’s epitope in more detail, recombinant OspC_A_ was subjected to HX-MS in the absence and presence of mouse B5 IgG. HX-MS provides peptide level resolution of antibody-antigen interactions in solution based on differential hydrogen-deuterium exchange rates between unbound and bound targets (Angalakurthi et al., 2018; Brier et al., 2021; Chen et al., 2019; Grauslund et al., 2021; Hodge et al., 2022; Malito et al., 2013; Seow et al., 2022; Vinciauskaite and Masson, 2022). We recently used HX-MS to localize a dozen human B cell epitopes the *B. burgdorferi* antigen, OspA (Haque et al., 2022). In the case of OspC, we first generated a peptidic map of OspC_A_, which yielded 87 peptides that covered the entire length of the molecule (**Figure S3**). By HX-MS, the OspC N- and C-terminal regions displayed a high degree of flexibility in the unbound state, as evidenced by a high degree of exchange (**data not shown**). The addition of B5 IgG resulted in weak, but statistically significant protection across the majority of the OspC molecule, possibly reflecting a combination of allosteric effects and dynamic interconversion between bound and unbound states of the antigen. The strongest protection, however, encompassed OspC_A_’s terminal α-helix 6, corresponding to peptidic residues 163-168, 171-172, 174-175, 177-179, 181-182, 184-186, 188-197, and 199-200 (**Figure 2**). These results suggest that binding of B5 IgG influences flexibility of the entire length of OspC’s terminal α-helix.

**Figure 2.**
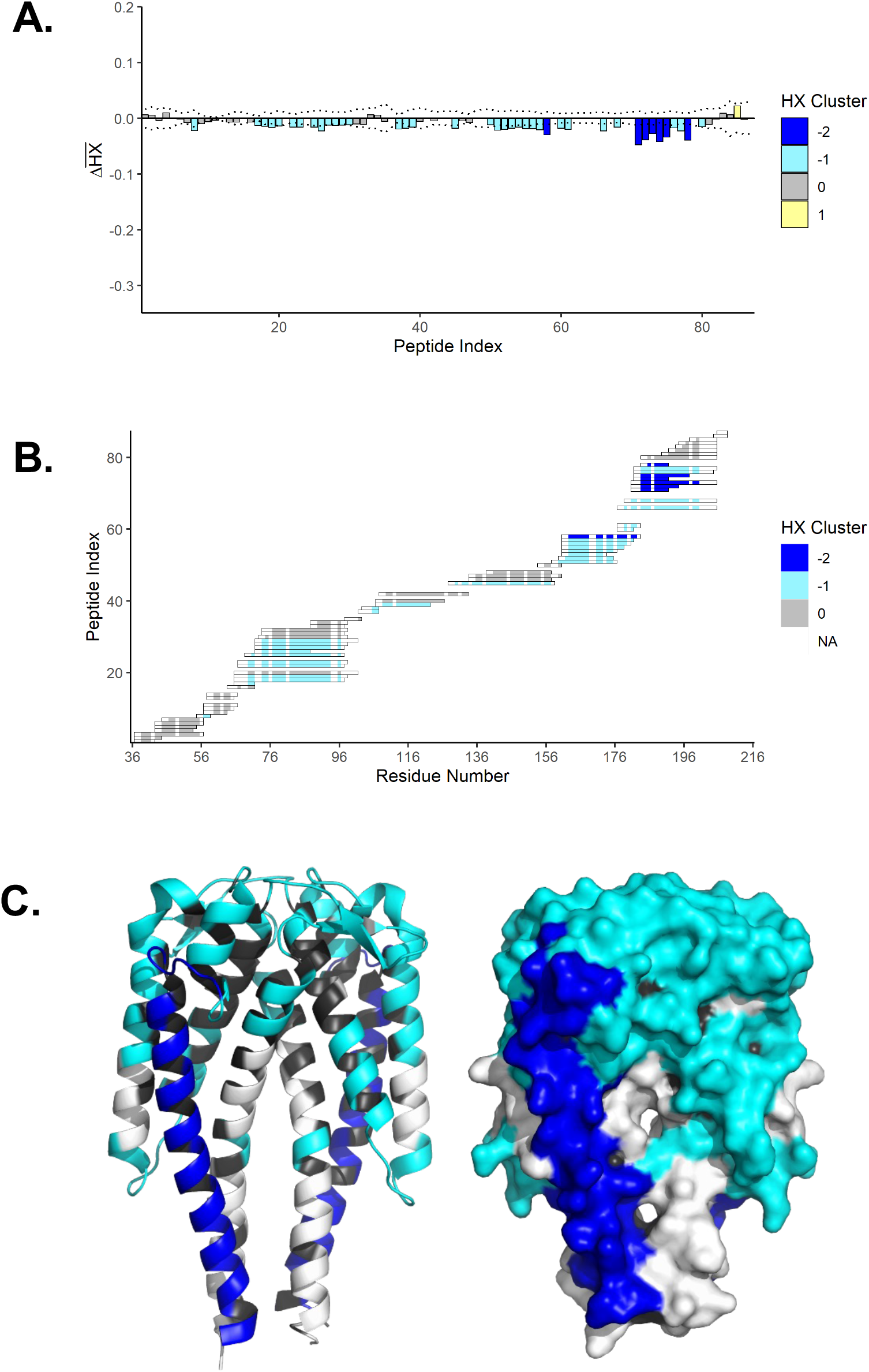
Localization of B5 IgG binding sites on OspC_A_ by HX-MS. (**A**) Time-averaged, normalized HX differences (Δ*HX*) between OspC_A_ and OspC_A_ bound to B5 IgG. Negative bars denote slower HX in the presence of B5 IgG. The peptide index organizes the peptides from N- to C-terminus. The values are color-coded based on k-means clustering. Extreme cluster values (e.g. −2) denotes stronger effects; (**B**) Peptide resolution map of the B5 IgG epitope showing clustered HX; (**C**) Cartoon (left) and surface (right) representations of OspC_A_ [PDB ID 1GGQ] with relative effects of B5 IgG as interpreted from panels A and B. The dark blue shading represents strongly protected regions of OspC_A_, light blue represents weak (but significant) protection, and black denotes lack of coverage.

### X-ray crystal structure of Fab B5-OspC_A_

To define B5’s epitope in greater detail, we solved the X-ray crystal structure of the B5 Fab fragment in complex with OspC_A_ at 2.7 Å resolution in the P2_1_2_1_2 space group. The crystal structure revealed two B5 Fabs bound to a single OspC_A_ homodimer (1:1 Fab:OspC_A_ stoichiometry) in a side-on fashion (**Figure 3**). The B5 Fab fragments (Fab, Fab’) made nearly identical contacts on opposite sides of the OspC_A_ homodimer, as described in detail below. B5 Fabs assumed a canonical structure with two heavy chain immunoglobulin domains (V_H_, C_H_1) and two light immunoglobulin domains (V_L_, C_L_) each containing 7-10 β-strands arranged in two β-sheets that folded into a two-layer sandwich with all six CDRs (L1-3, H1-3) on one face of the molecule. The homodimeric structure of OspC_A_ unbound [PDB 1GGQ] and bound to B5 Fabs were similar, as evidenced by a Root-Mean-Square Deviation (RMSD) of 1.0 Å. Thus, antibody engagement did not induce any major conformational changes OspC_A_.

**Figure 3.**
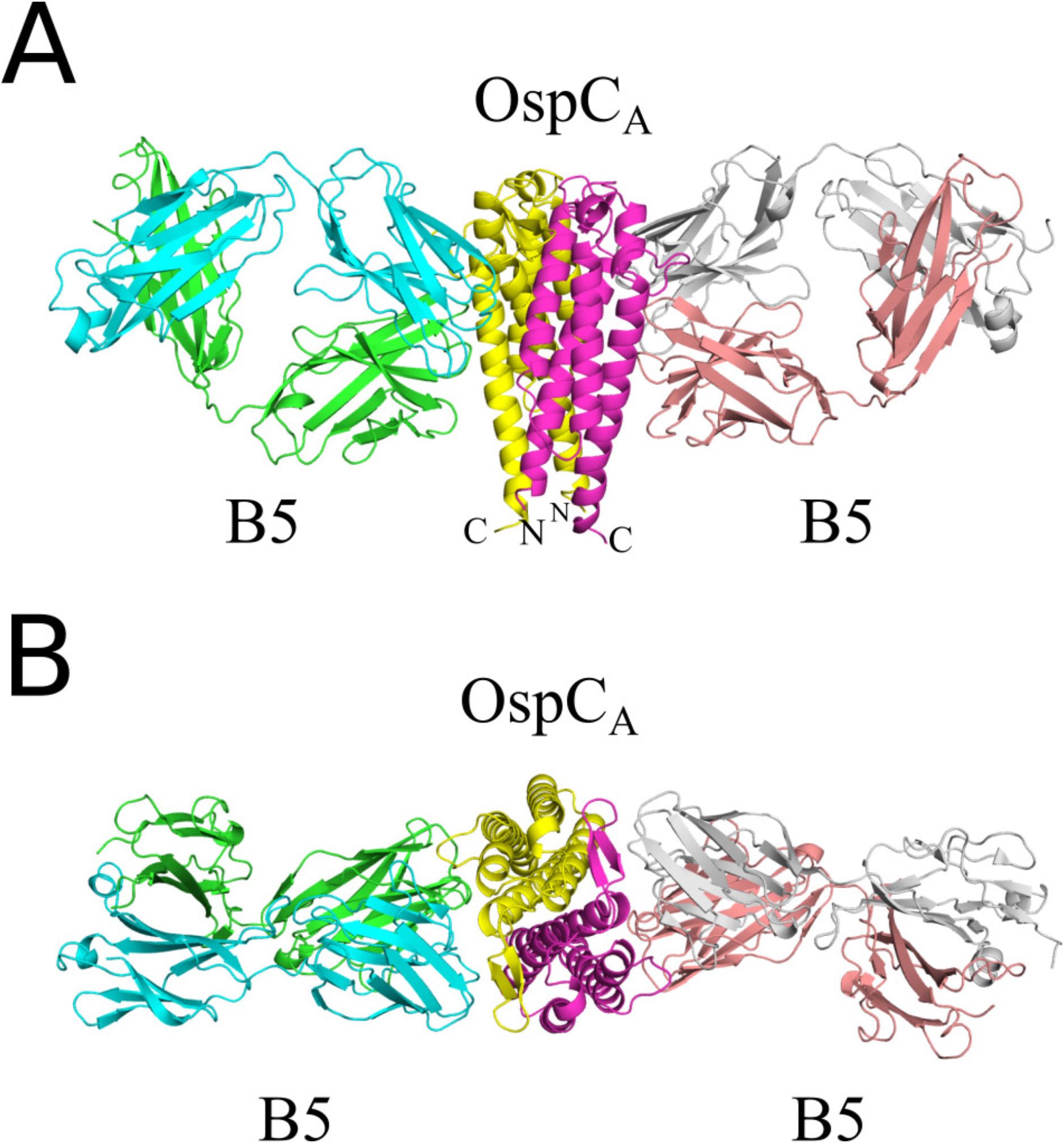
Structure of B5 Fab-OspC_A_. (A) Side-on and (B) top-down ribbon diagrams of OspC_A_ homodimer (OspC_A_, OspC_A_’) in complex with B5 Fabs (B5, B5’). The OspC_A_ is colored in yellow and OspC_A_’ in magenta. The B5 Fab V_H_ and C_H_1 elements are colored light green and the V_L_ and C_L_ cyan. The B5’ Fab V_H_ and C_H_1 elements are colored in salmon red and V_L_ and C_L_ in light gray. The N- and C-termini of OspC_A_ and OspC_A_’ are labelled accordingly (N, C).

The interaction between B5 Fab and OspC_A_ buried a total surface area of 2,040 Å^2^ (2,002 Å^2^ for the second B5-OspC_A_ interface within the asymmetric unit) establishing nine hydrogen bonds and three salt bridges (12 hydrogen bonds and three salt bridges for the second B5-OspC_A_ interface within the asymmetric unit) (**Table 2**; **Figure 4**). The B5 Fab V_H_ domain contributed slightly more to the interaction burying 1042 Å^2^ (1027 Å^2^ for the second B5-OspC_A_ contact within the asymmetric unit) than the V_L_ domain which buried 998 Å^2^ (975 Å^2^ for the second B5-OspC_A_ interface in the asymmetric unit). The B5 V_H_ domain formed four hydrogen bonds (three hydrogen bonds for the second B5-OspC_A_ interface in the asymmetric unit) that includes CDR-H1 residue Tyr-27 with OspC_A_ Lys-175, and CDR-H3 residue Tyr-102 and OspC_A_ Glu-71 (**Figure 4**). The B5 V_H_ domain also accounted for two of the three salt bridges observed between B5 and OspC_A_. The two salt bridges formed between B5’s V_H_ domain and OpsCA involved H2 Glu-54 with OspC_A_’s Lys-188 and H1 His-32 with Glu-181. The third salt bridge involved V_H_ framework residue Glu-1 and OspC_A_ Lys-166 (**Figure 4**). The B5 V_L_ domain formed five hydrogen bonds (eight hydrogen bonds for the second B5-OspC_A_ interface within the asymmetric unit) including CDR L2 Thr-52 with OspC_A_ Lys-79, and CDR L2 Tyr-49 with OspC_A_ Thr-162. There were no salt bridges between B5 V_L_ domain and OspC_A_.

**Figure 4.**
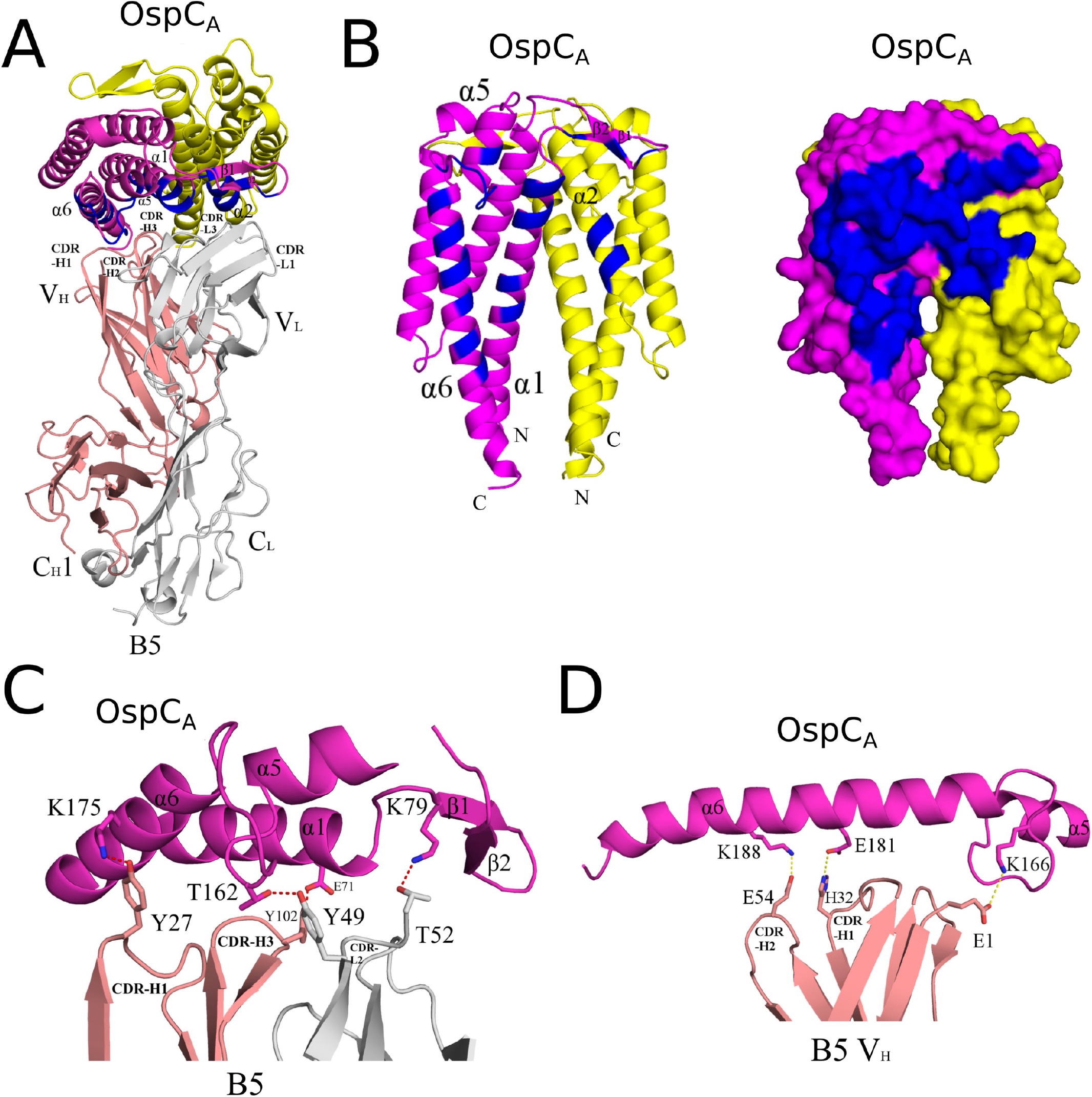
Detailed interactions between B5 and OspC_A_ revealed from the co-crystal structure. (**A**) Ribbon structure (top-down view) of OspC_A_ homodimer (OspC_A_, magenta; OspC_A_’, yellow) in complex with a single B5 Fab (V_H_ and C_H_1 elements, salmon red; V_L_ and C_L_, light gray). The OspC_A_ residues that engage with B5 are colored blue. Key secondary structures are labeled (a-helices 1, 2, 5, and 6; β-strands 1 and 2); (**B**) Ribbon (left) and surface (right) depiction of an OspC_A_ homodimer (OspC_A_, magenta; OspC_A_’, yellow) with B5 interacting residues shaded in dark blue. OspC_A_ N and C-termini are labelled N and C, respectively. Representations of key (**C**) H-bonds (red dashes) and (**D**) salt bridges (yellow dashes) between OspC_A_ (magenta) and Fab B5 (salmon red in panel C; gray in panel D). Side chains are drawn as sticks and color coordinated to the main chain color, with nitrogen atoms shaded blue and oxygen atoms shaded red. CDR elements are labelled per convention: CDR-L1, -L2, -L3; CDR-H1, -H2, -H3.

**Table 2.**
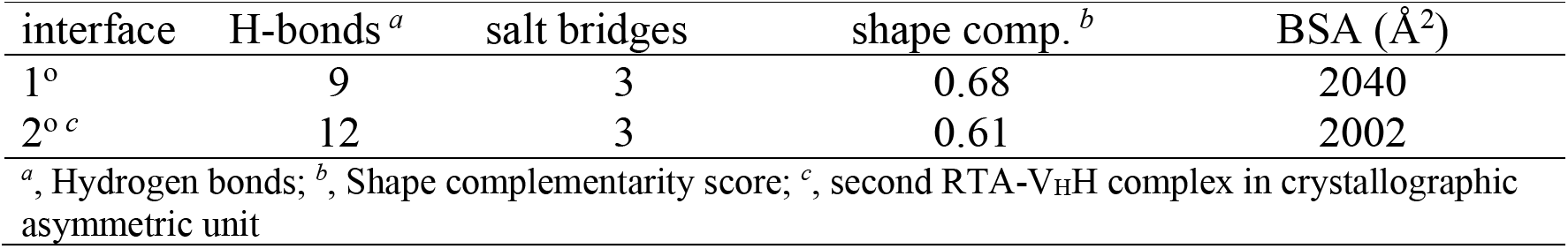
Summary of OspC_A_-B5 binding data and interface information.

Collectively, the B5 CDR H1, H2, and H3 elements contacted OspC_A_ α-helix 1 and α-helix 6, along with the loop between a-helices 5 and 6 (loop 5-6). The CDR H3 element also buried 30 Å^2^ of α-helix 2’ in the absence of any H-bonds or salt bridges. CDRs L1, L2, and L3 contacted OspC_A_ a-helices 1 and 5, β-strands 1 and 2, as well as the loop region immediately N-terminal to α-helix 6. CDR L1 and L3 interacted with α-helix 2’, burying 364 Å^2^ and forming a single H-bond between CDR L1 Asn-30 and Gln-110 in OspC_A_. The fact that B5 Fabs straddle the OspC dimer interface not only explains conformation-dependent nature of B5’s epitope, but more broadly the observation that immunizing mice with homodimeric OspC is more effective than monomeric OspC at eliciting protective antibodies (Edmondson et al., 2017).

### Structural basis of B5 IgG specificity for OspC_A_

To elucidate the structural basis for B5 IgG’s specificity for OspC_A_, we solved the crystal structures of recombinant OspC_B_ and OspC_K_ at 1.5 Å and 1.9 Å resolution, respectively, in the P2_1_ space group (**Figure S4**). OspC_B_ and OspC_K_ each formed homodimers nearly identical to OspC_A_ (unbound or bound to B5). Specifically, OspC_B_ and OspC_K_ monomers each consisted of four long a-helices (1, 2, 3, 6), two shorter a-helices (4 and 5) and a two stranded β-sheet (**Figure S4**). The RMSD between the homodimeric OspC_A_ (bound to B5) versus OspC_B_ and OspC_K_ ranged from 0.8 Å to 1.4 Å. In each case, the OspC dimer interface is predominantly hydrophobic, with ~80% of the protein atoms in the interface being nonpolar. The monomeric form of OspC_A_ (with or without B5 Fab bound) was structurally more similar to OspC_K_ than OspC_B_, with an RMSD of ~0.7 Å. A deletion at residue 74 and an insertion of residue 165 in OspC_B_ relative to OspC_A_ and OspC_K_ accounts for its significantly greater structural deviation, as exhibited by an RMSD of 0.9 Å when superposed onto OspC_A_ or OspC_K_ monomers. After molecular replacement calculations were performed, the resulting phase information was used to calculate electron density maps utilized to manually insert the correct residues into each model and manually build other regions of each model for the OspC_B_ and OspC_K_ structures. Crystallographic and refinement data for each structure demonstrated a refined molecular model with excellent agreement to the crystallographic data, as well as excellent geometry (**Table S1**).

Superpositioning the B5-OspC_A_ complex onto OspC_B_ and OspC_K_ revealed several structural attributes that likely account for B5’s inability to recognize OspC_B_ and OspC_K_. One prominent feature involves the contact between B5 Trp-100 with Gly-174 in OspC_A_. In OspC_B_ and OspC_K_, Glu-175 is superposed with OspC_A_ Gly-174. The bulkier glutamic acid (Glu) side chain at position 175 would be expected to sterically clash with Trp-100 (**Figure 5**; **Figure S5**). Specifically, the side chain positions of Trp-100 (B5) and Glu-175 (OspC_B_, OspC_K_) are both highly preferred rotamers that would impede a B5 interaction. Furthermore, in the case of OspC_B_, an “insertion” of Ala-165 within loop 5-6 in OspC_B_ relative to OspC_A_ alters the configuration of the loop resulting in a theoretical clash with Tyr-49 of B5 (**Figure 5B**; **Figure S5**). OspC_B_ is also incapable H-bonding with B5 Tyr-49, due to an Ala rather than a Thr at position 162, as is the case in OspC_A_. The absence of this H-bond donor would be expected to compromise the B5-OspC_B_ interaction. Another structural feature in OspC_B_ disfavoring B5 binding includes deletion of one residue immediately before Lys-74 in OspC_B_. This deletion substantially altered residue positions 73-75 within α-helix 1 of OspC_B_ relative to the same residues in OspC_A_ and OspC_K_. As a result, the superposed side chain of Lys-74 in OspC_B_ clashes with B5 Tyr-102. Though several preferred rotamers of OspC_B_ Lys-74 can readily pivot away from B5 Tyr-102 alleviating the close encounter between these two residues, movement of Lys-74 away from Tyr-102 and B5 residue Ala-50 would significantly reduce contact between B5 and OspC_B_ by ~50 Å^2^ likely diminishing B5 binding to OspC_B_ (**Figure 5C**; **Figure S5**). In the case of OspC_K_, divergent primary amino acid sequences at consequential residues certainly contribute to lack of B5 recognition. For example, OspC_A_ Lys161 forms a π-cation interaction with B5 Trp-100, which cannot occur in the case of OspC_K_ due to an Ile residue at this position (**Figure 5D**; **Figure S5**). To determine the potential cross-reactivity of B5 with the other OspC types, we looked at the primary sequence conservation across each OspC focusing on regions in OspC_A_ that support B5 binding while also considering regions in OspC_B_ and OspC_K_ that seemingly antagonize B5 interaction. Two two OspC types, C3 and I3, with 76% and 78% overall sequence identities to OspC_A_, would predictably interact with B5. C3 and I3 possess similar sequences within α-helix 1 and loop 5-6 along with a few other key B5 contact residues found in OspC_A_ (**Figure S5**). For example, OspC_13_ is similar to OspC_A_ in that it has a Gly at position 174, providing ample room to contact B5’s Trp-100. In the case of OspCc3, an Asp residue replaces Gly-174; while the Asp side chain is larger than Gly there is still sufficient space to engage Trp-100 in B5. It is interesting that Baum and colleagues demonstrated that OspC_A_ and OspC_13_ were most immunologically cross-reactive pairs in a protein microarray consisting of 23 OspC types (Baum *et al*., 2013). Elucidating structural insights into the molecular interactions that promote or repel protective antibodies has important implications for rational vaccine design.

**Figure 5.**
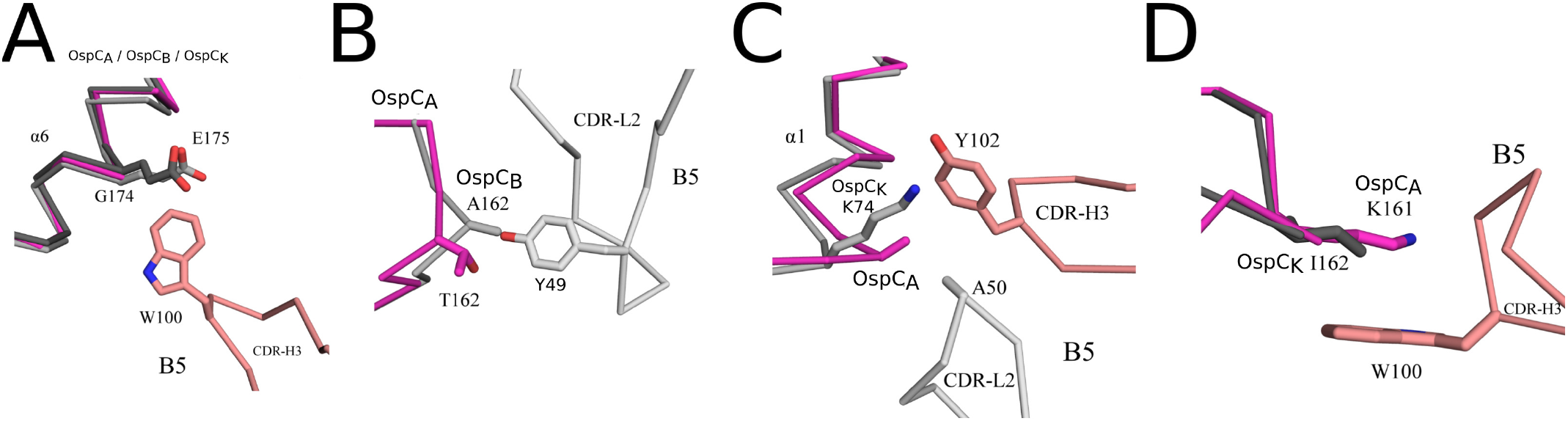
Structural basis of B5 specificity for OspC_A_. Interface between B5 Fab and OspC_A_ superposed with OspC_B_ and OspC_K_, highlighting interactions that are conceivably disrupted by sequence deviations among OspC types. (**A**) The C-a traces of OspC_A_ (magenta) bound to B5 (salmon red) with superposed OspC_B_ (light gray) and OspC_K_ (dark gray) highlighting potential steric clash between Glu-175 of OspC_B_ and OspC_K_ with Trp-100 of B5 Fab. (**B**) C-a traces of OspC_A_ (magenta) bound to B5 (salmon red) superposed with OspC_B_ (light gray). The image highlights likely repulsion between Ala-162 of OspC_B_ with Tyr-49 of B5 Fab. (**C**) C-a traces of OspC_A_ (magenta) bound to B5 with CDR-H3 (gray) and CDR-L2 (gray) superposed with OspC_K_ (light gray). The image highlights a potential clash between Lys-74 of OspC_K_ and B5’s Tyr-102. (**D**) The C-α trace of OspC_A_ (magenta) bound to B5 (salmon red) superposed with OspC_K_ (dark gray). Ile-162 in OspC_K_ precludes a potential π-cation interaction that occurs between OspC_A_ Lys-161 and Trp-100 in B5 Fab. Side chains are drawn as sticks and color coordinated to the main chain color, with nitrogen atoms shaded blue and oxygen atoms shaded red.

### Overlap between B5’s footprint and human linear B cell epitopes on OspC

Numerous studies have identified OspC-derived peptides that react with human sera from Lyme disease patients and that have been proposed to have utility of serodiagnosis (**Table 3**). Positioning of seven such linear epitopes on the structure of OspC_A_ revealed that four of them fall within B5’s footprint (**Figure 6**). These results reveal overlap between conformation-dependent and -independent epitopes on the surface of OspC.

**Figure 6.**
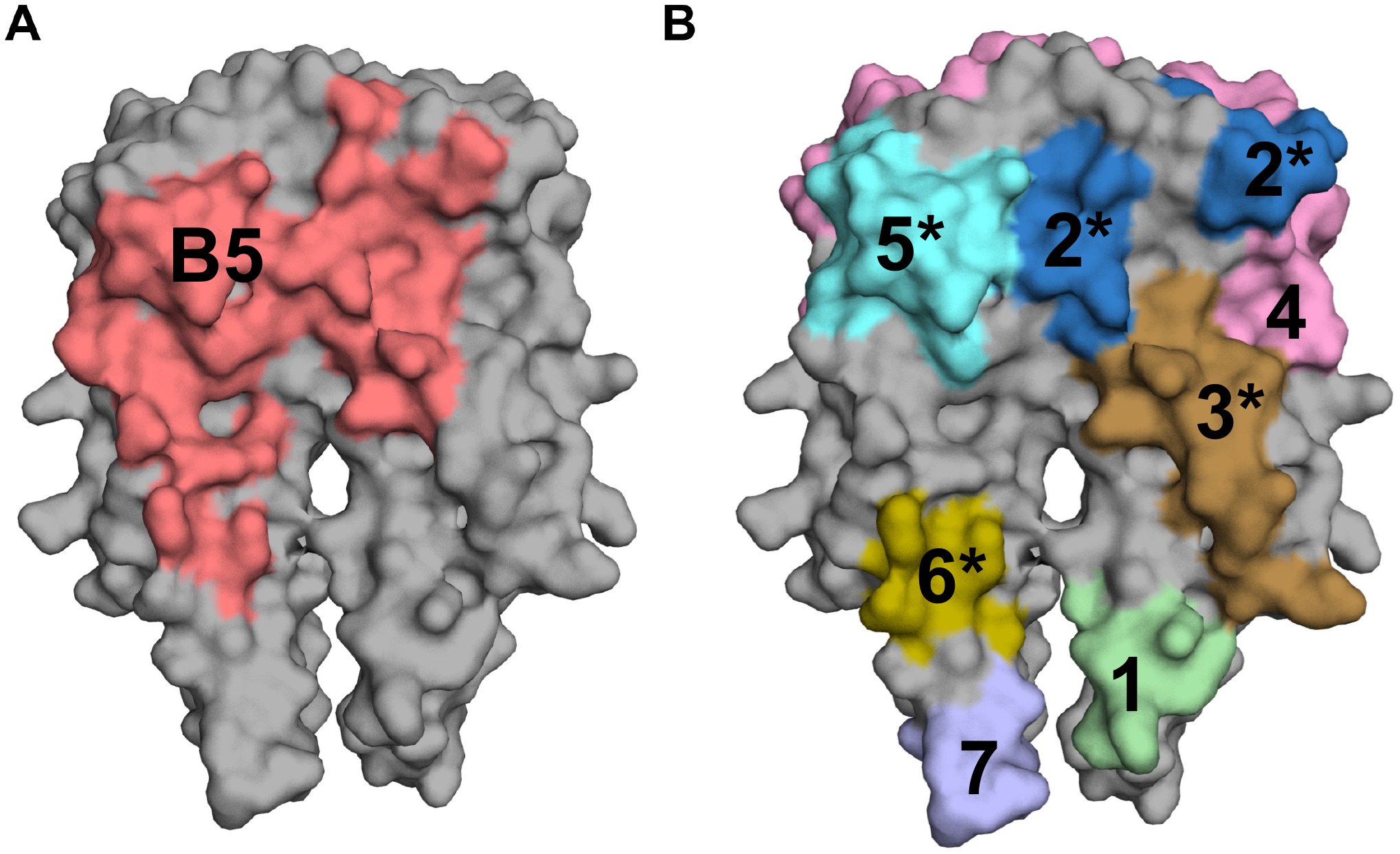
Depiction of overlap between linear human B cell epitopes and B5 contact points on OspC. Surface representation of a OspC_A_ homodimer [shaded gray; PDB ID 1GGQ] with (**A**) B5 contact points (salmon red) from **Figure 3** and (**B**) the location of seven (1-7) previously reported linear B cell epitopes on OspC and reported in **Table 3**. The asterisks (*) indicate proposed overlap with B5 contact points.

**Table 3.**
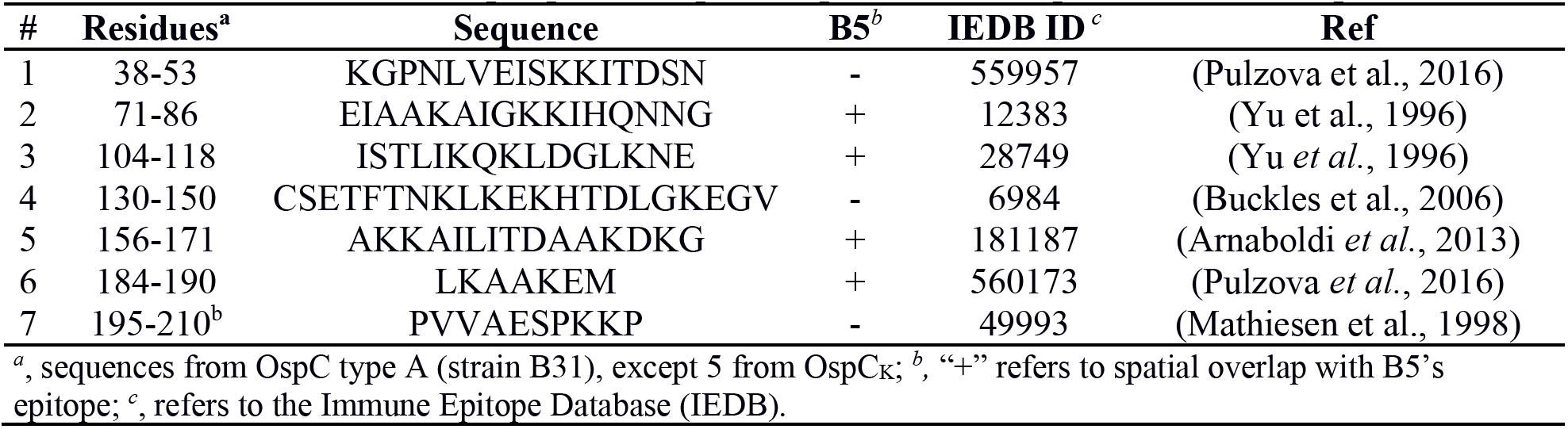
Human linear B cell epitopes on OspC and putative overlap with B5 contact points.

### Conclusion

OspC is a multifunctional lipoprotein that facilitates both *B. burgdorferi* tick-to-host transmission and infectivity (Caine and Coburn, 2016). OspC is also of interest as a next generation Lyme disease vaccine (Dattwyler and Gomes-Solecki, 2022; Wormser, 2022). Although structures of OspC types A, E, and I were solved two decades ago, the identification of protective B cell epitopes on OspC has been limited to linear (peptide) targets (Arnaboldi et al., 2013; Earnhart *et al*., 2005; Norek and Janda, 2017). Thus, the structure reported here between OspC_A_ and Fabs derived from the mouse MAb, B5, represents the first high-resolution image of a protective B cell epitope on OspC. The B5 Fab-OspC_A_ complex is stunning in that B5 attacks OspC in a side-on fashion and contacts residues (Asp-70, Glu-71) that form a cavity proposed to serve as binding site for tick- or mammalian-derived ligands (Eicken *et al*., 2001; Kumaran *et al*., 2001). The structure also revealed that B5 Fabs straddle the OspC_A_-OspC_A_’ dimer interface, thereby explaining (at least in part) the conformation-dependent nature of B5’s epitope (Gilmore and Mbow, 1999). Ultimately the results of this study advance our understanding of serotype-specific immunity to *B. burgdorferi*, as well as opening avenues for structure-based OspC vaccine development (Ward and Wilson, 2020).

## Materials and Methods

### Mouse and chimeric B5 IgG MAbs

Lyophilized B5 IgG from CDC was reconstituted to final concentration of 6 mg/ml. B5 Fab fragments were generated by papain digestion followed by affinity depletion of the Fc fragment by Protein A FPLC chromatography. The resulting B5 Fab was purified to homogeneity by size-exclusion chromatography (SEC) using a Superdex 200 16/60 gel filtration column. The B5 mouse hybridoma was cultured as described from frozen aliquot (Mbow *et al*., 1999). In addition, to ensure sufficient supply of B5 MAb, the mouse B5 V_H_ and V_L_ regions were cloned into human IgG1 Fc and k light chain expression vectors and used to transfect Expi293 cells following protocols previously described (Wang et al., 2016). The resulting chimeric B5 IgG1 was purified and used for dot blot and flow cytometry analysis. Fabs were generated from murine B5 IgG by papain digestion followed by affinity depletion of the Fc fragment by Protein A FPLC chromatography. The resulting B5 Fab was purified to homogeneity by size-exclusion chromatography (SEC) using a Superdex 200 16/60 gel filtration column.

### Affinity measurement using biolayer interferometry (BLI)

Biotinylated OspC_A_ (3 μg/mL) in buffer (PBS containing 2% w/v BSA) was captured onto streptavidin biosensors (#18-5019, Sartorius, Goettingen, Germany) for 5 min. After 3 min of baseline in buffer, sensors were then exposed to a 2-fold serial dilution of MAb B5, ranging from 100 to 1.56 nM, for 5 min. The sensors were then dipped into wells containing buffer alone for 30 min. An eighth sensor was also loaded with biotinylated-OspC_A_, but not exposed to MAb B5, and was thus used as a background drift control, and subtracted from the other sensor data. The raw sensor data was then loaded into the Data Analysis HT 12.0 software, and the data was fit to a 1:2 bivalent analyte model. Data was captured on an Octet RED96e Biolayer Interferometer (Sartorius) using the Data Acquisition 12.0 software.

### Indirect fluorescent antibody staining and flow cytometry

A derivative of *B. burgdorferi* strain B31 over expressing OspC was kindly provided by Dr. Yi-Pin Lin (Wadsworth Center). The strain was cultured in BSK-II media at 37°C with 5% CO2 to mid-log phase. Cells were collected by centrifugation (3,300 x *g*), washed with PBS, resuspended in BSK-II (minus phenol red indicator) at a final concentration of 1×10^8^ cells/ml, and incubated at room temperature for 30 min. A total of 5×10^6^ cells in 50 μl were incubated with 10 μg/ml of chimeric B5 IgG1 at 37°C for 1 h. Incubation with an isotype control, PB10, was included as a negative control (Rong et al., 2020). The reaction volume was then increased with the addition of 450 μl of BSK-II (minus phenol red), and 2° ab (goat anti-human IgG (H+L)-Alexa 647; Invitrogen) was added at 1/500 dilution and allowed to incubate at 37°C for 30 min. Alexa-647 labeled cells were analyzed on a BD FACS Calibur flow cytometer. Data was obtained and analyzed using BD’s CellQuest Pro software.

### Dot blot

Recombinant OspC types A, B, and K (**Table 1**) at 0.5 μg/ul, were 5-fold serially diluted in PBS, and 2 μl of each dilution was spotted on a dry nitrocellulose membrane. The spots were allowed to air-dry for 1 h, then blocked with 5% milk in 1x TBS-T for 18 h, and incubated with 0.1 μg/ml chimeric B5 IgG1 in 5% milk 1x TBS-T at room temperature for 1 h. The membrane was then washed twice with 1x TBS-T, incubated with a 1:10,000 dilution goat anti-human IgG (H+L)-HRP (Invitrogen), and washed twice more before detection with ECL (ECL Plus Western Blotting Substrate; Pierce, ThermoFisher, Waltham, MA). Images were acquired and analyzed using an iBright 1500 (Invitrogen).

### OspC ELISA

B5 IgG was coated onto wells of a 96 well Immulon 4HBX plate (ThermoFisher, Waltham, MA) at 1 μg/mL in PBS overnight at 4°C. Wells were then blocked for 2 h at room temperature with 2% goat serum in 0.1% Tween-20 in PBS. Biotinylated OspC Types A, B and K were then diluted 2-fold across the plate, starting at 20 μg/mL. Plates were washed, and then captured biotinylated OspC was detected with avidin-HRP (Pierce, Rockford, IL) for 1 h. Plates were washed again, and capture was visualized with SureBlue TMB (SeraCare, Milford, MA). The reaction was stopped with 1M phosphoric acid, and the optical density at 450 nm was read using a SpectraMax iD3 (Molecular Devices, San Jose, CA).

### LC-MS analysis

For peptic peptide mapping, recombinant OspC_A_ was diluted with quench solution (200 mM glycine, pH 2.5) and 50 pmol sample was injected in each run. OspC_A_ was digested by in-house prepared immobilized pepsin column (2.1×50 mm) (Wang et al., 2002). Digested peptides were by trapped and desalted by C-8 (Zorbax 300SB C8, 2.1 x 12.5 mm, 5 μm particles) for 120 s and separated by a C-18 column (Zorbax 300SB 2.1 × 50 mm, 3.5 μm particle diameter, Agilent, Santa Clara, CA). For LC, mobile phase A was 0.1% formic acid in water, and B was 0.1% formic acid in acetonitrile. A total of 25 min LC method, 10 min with 15%-35%B was used to separate peptides and 15 min was used for cleaning purpose. Peptide were detected and mass was measured by quadruple time of flight mass spectrometer Q-TOF mass spectrometer (Agilent 6530 in ESI-positive ion mode). All the peptic peptides were assigned by tandem mass spectrometry (collision induced dissociation (CID) fragmentation). Agilent MassHunter Qualitative Analysis with BioConfirm (version B.07.00) software was used for the analysis of all the mass spectrometry data. A total of 87 peptides were identified and mapped in Fig. S1. This map shows 100% OspC_A_ sequence coverage with median length 17.0 residues and 8.6 average redundancy.

### Hydrogen exchange-mass spectrometry (HX-MS) and data analysis

OspC_A_ and B5 MAb were buffer exchanged and assayed by our previous protocol (Haque *et al*., 2022). Previously flash frozen OspC_A_ (19μM) was thawed at room temperature. B5 sample was collated from 4°C and buffer exchanged on the day of experiment. Free protein state (OspC_A_) sample was prepared by diluting 19μM OspC_A_ to 9μM by the addition of 20 mM phosphate, 100 mM NaCl, pH 7.40. Bound state (OspC_A_ +B5) was prepared by three strokes mixing and adjusted to a final concentration of 9 μM. HX-MS labeling conditions, robot methods, maximally deuterated experiment (OspA paper) protocol, data analysis was done as recently reported (Haque *et al*., 2022).

### Cloning, expression, and purification of OspC

The PCR amplicon encoding *B. burgdorferi* OspC_A_ (residues 38 to 201) was subcloned into the pSUMO expression vector that contained an N-terminal deca-histidine and SUMO tag. The PCR amplicons for *B. burgdorferi* OspC_B_ and OspC_K_ containing residues 38 to 202 were subcloned into the pMCSG7 expression vector that contained an N-terminal deca-histidine tag. Cloning was performed using standard ligase independent cloning (LIC). All OspC types were expressed in *E. coli* strain BL21 (DE3). The transformed bacteria were grown at 37°C in TB medium and induced at 20°C at an OD_600_ of 0.6 with 0.1 mM IPTG for ~16 h. After induction, cells were harvested and resuspended in 20 mM HEPES [pH 7.5] and 150 mM NaCl. The cell suspension was sonicated and centrifuged at 30,000 x *g* for 30 min. After centrifugation, the protein-containing supernatant was purified by nickel-affinity and size-exclusion chromatography on an AKTAxpress system (GE Healthcare), which consisted of a 1 mL nickel affinity column followed by a Superdex 200 16/60 gel filtration column. The elution buffer consisted of 0.5 M imidazole in binding buffer, and the gel filtration buffer consisted of 20 mM HEPES [pH 7.5], 150 mM NaCl, and 20 mM imidazole. Fractions containing each OspC type were pooled and subject to TEV protease cleavage (1:10 weight ratio) for 3 h at room temperature to remove respective fusion protein tags. The cleaved proteins were passed over a 1mL Ni-NTA agarose (Qiagen) gravity column to remove TEV protease, cleaved residues, and uncleaved fusion protein. After purification, Fab B5 was complexed with OspC_A_ in a 1:1 stoichiometry, then concentrated to 10 mg/ml final for all crystallization trials.

### Crystallization and data collection

All crystals were grown by sitting drop vapor diffusion using a protein to reservoir volume ratio of 1:1 with total drop volumes of 0.2 μl. Crystals of the B5 Fab-OspC_A_ complex were produced at 22°C using a crystallization solution containing 100 mM sodium cacodylate pH 6.5, 5% PEG 8K, and 40% MPD. Crystals of the OspC_B_ were produced at 22°C using a crystallization solution containing 100 mM sodium phosphate citrate pH 4.2, 41.9% PEG 600 Crystals of the OspC_K_ were produced at 4°C using a crystallization solution containing 100 mM Tris pH 8.5, 40% PEG 400, 200 mM LiSO4, 10 mM 2-aminoethanesulfonic acid. All crystals were flash frozen in liquid nitrogen after a short soak in the appropriate crystallization buffers supplemented with 25% ethylene glycol. Data were collected at the 24-ID-E beamline at the Advanced Photon Source, Argonne National Labs. All data was indexed, merged, and scaled using HKL2000 (Otwinowski and Minor, 1997) then converted to structure factor amplitudes using CCP4 (Winn et al., 2011).

### Structure determination and refinement

The B5 Fab-OspC_A_ complex, OspC_B_, and OspC_K_ structures were solved by molecular replacement using Phaser (Otwinowski and Minor, 1997). Molecular replacement calculations were performed using the coordinates of the murine monoclonal Fab 3E6 V_H_ and C_H_1 domains (PDB ID: 4KI5) along with the V_L_ and C_L_ domains of the human germline antibody hepatitis E virus E2S antibody (PDB ID: 3RKD) as the search model for Fab B5 in the B5-OspC_A_ complex. The OspC coordinates (PDB ID: 1GGQ) were used as the search model for OspC_A_ in the OspC_A_-B5 complex. The same OspC coordinates (PDB ID: 1GGQ) were also used as search models for the OspC_B_ and OspC_K_ structure determinations. The resulting phase information from molecular replacement was used for some manual model building of each structure solved using the graphics program COOT (Emsley et al., 2010) and structural refinement employing the PHENIX package (Adams et al., 2010). Data collection and refinement statistics are listed in **Table S1**, as are the Protein Data Bank (http://www.rcsb.org/pdb/) codes associated with each of the three structures generated in this study (B5-OspC_A_, PDB ID 7UIJ; OspC_B_, PDB ID 7UJ2; OspC_K_, PDB ID 7UJ6). Molecular graphics were prepared using PyMOL (Schrodinger, DeLano Scientific LLC, Palo Alto, CA).

## Acknowledgements

We gratefully acknowledge the Wadsworth Center’s Tissue and Cell culture facility for preparing BSK II media, and Dr. Renjie Song in the Immunology Core for assistance with flow cytometry. We thank Dr. Timothy LaRocca (Albany College of Pharmacy and Health Sciences) and Dr. Graham Willsey for insightful discussions and assistance. We thank Elizabeth Cavosie for administrative assistance. This work was supported by the National Institute of Allergy and Infectious Diseases (NIAID), National Institutes of Health, Department of Health and Human Services Contract No. 75N93019C00040. This work is based upon research conducted at the Northeastern Collaborative Access Team beamlines, which are funded by the National Institute of General Medical Sciences from the National Institutes of Health (P30 GM124165). The Eiger 16M detector on the 24-ID-E beam line is funded by a NIH-ORIP HEI grant (S10OD021527).

---

**Table S1.** Data associated with crystal structures reported in this study.

|  |  |  |  |
| --- | --- | --- | --- |
| **Data Collection** |  |  |  |
| Complex | OspC<sub>A</sub>-B5 | OspC<sub>B</sub> | OspC<sub>K</sub> |
| Space group | P2<sub>1</sub>2<sub>1</sub>2 | P2<sub>1</sub> | P2<sub>1</sub> |
| Cell parameters: *a, b, c*<br>(Å) / β (°) | 54.7 / 139.8 / 205.7 | 42.9 / 44.1 / 68.8 /<br>103.8 | 52.3 / 52.3 / 116.7 /<br>92.5 |
| BNL Beamline | 21-ID-E | 21-ID-E | 21-ID-E |
| Resolution range <sup>a</sup>(Å) | 50-2.70 (2.75-2.70) | 50-1.50 (1.53-1.50) | 50-1.95 (1.98-1.95) |
| wavelength (Å) | 0.979 | 0.979 | 0.979 |
| No. of reflections | 3988365 | 3312121 | 2685656 |
| Average redundancy<sup>a</sup> | 2.8 (2.7) | 2.7 (2.4) | 2.8 (2.6) |
| (*I*)/(*δ*)<sup>a</sup> | 21.1 (1.2) | 31.5 (3.2) | 17.6 (1.7) |
| Completeness<sup>a</sup> (%) | 99.6 (99.6) | 95.8 (95.6) | 97.8 (99.2) |
| *R*<sub>merge</sub> <sup>a,b</sup> (%) | 8.3 (145.0) | 5.9 (32.5) | 14.9 (131.1) |
| CC<sub>1/2</sub> <sup>a,c</sup> | (0.50) | (0.93) | (0.54) |
| **Refinement** |  |  |  |
| Bragg spacings <sup>a</sup>(Å) | 49.4-2.7 (2.76-2.70) | 36.8-1.5 (1.54-1.50) | 39.7-1.95 (1.99-1.95) |
| *R*<sup>d</sup> / *R*<sub>free</sub> <sup>e</sup> (%) | 20.4 / 25.6 | 14.7 / 16.7 | 19.3 / 22.8 |
| No. of Protein atoms | 8825 | 4908 | 4708 |
| No. of Waters | 32 | 270 | 358 |
| RMSD bond length (Å) | 0.005 | 0.012 | 0.003 |
| RMSD bond angle (°) | 0.78 | 1.14 | 0.52 |
| Ramachandran<br>favored / allowed<sup>f</sup> (%) | 94.8 / 99.8 | 97.4 / 100 | 97.0 / 100 |
| PDB code | **7UIJ** | **7UJ2** | **7UJ6** |
| <sup>a</sup> Values in outermost shell are given in parentheses.<br><sup>b</sup> $R_{\text{merge}} = (\sum \vert I_i - \langle I_i \rangle\vert ) / \sum \vert I_i\vert $, where *I<sub>i</sub>* is the integrated intensity of a given reflection.<br><sup>c</sup> $CC_{1/2} = (1 + q^2 \sigma_e^2 / \langle I \rangle 2^{-1})^{-1}$, where $\sigma_e$ denotes the mean error within a half-dataset, CC<sub>1/2</sub> is the correlation coefficient of two split data sets each derived by averaging half of the observations for a given reflection.<br><sup>d</sup> $R = \sum \vert \vert F_o\vert - \vert F_c\vert \vert / \sum \vert \vert F_o\vert \vert $, where *F<sub>o</sub>* and *F<sub>c</sub>* denote observe and calculated structure factors, respectively.<br><sup>e</sup> *R*<sub>free</sub> was calculated using 5% of data excluded from refinement.<br><sup>f</sup> Calculated using Molprobit. | | | |

**Figure S1.**
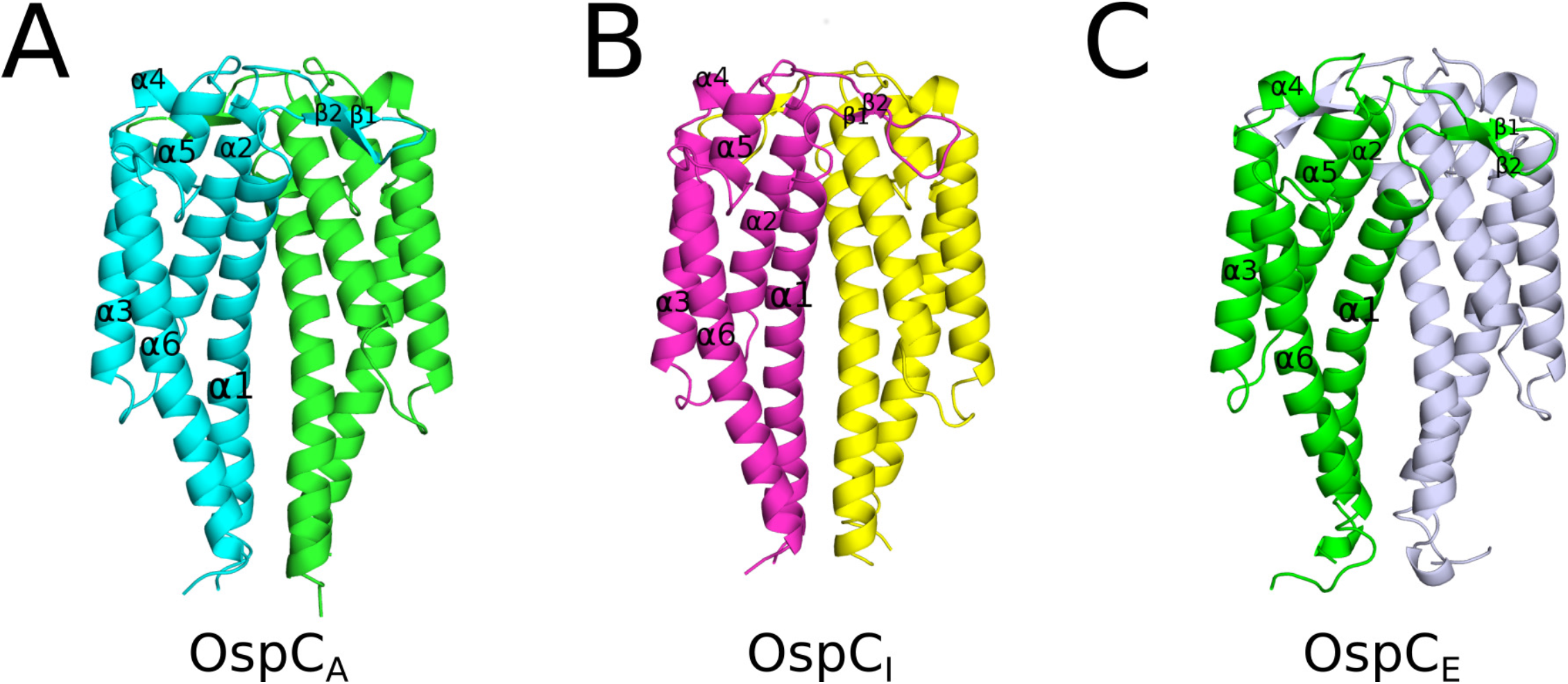
Structures of OspC types A, I and E. (A) Previously reported OspC structures. Ribbon diagrams of (A) OspC_A_ [PDB ID **1GGQ**] colored green and cyan,; (B) OspC_I_ [PDB ID **1F1M**] colored yellow and magenta; (C) OspC_E_ [PDB ID **1G5Z**] colored light blue and dark green. All a-helices (1-6) and β-strands 1 and 2 are labelled highlighting structural similarity.

**Figure S2.**
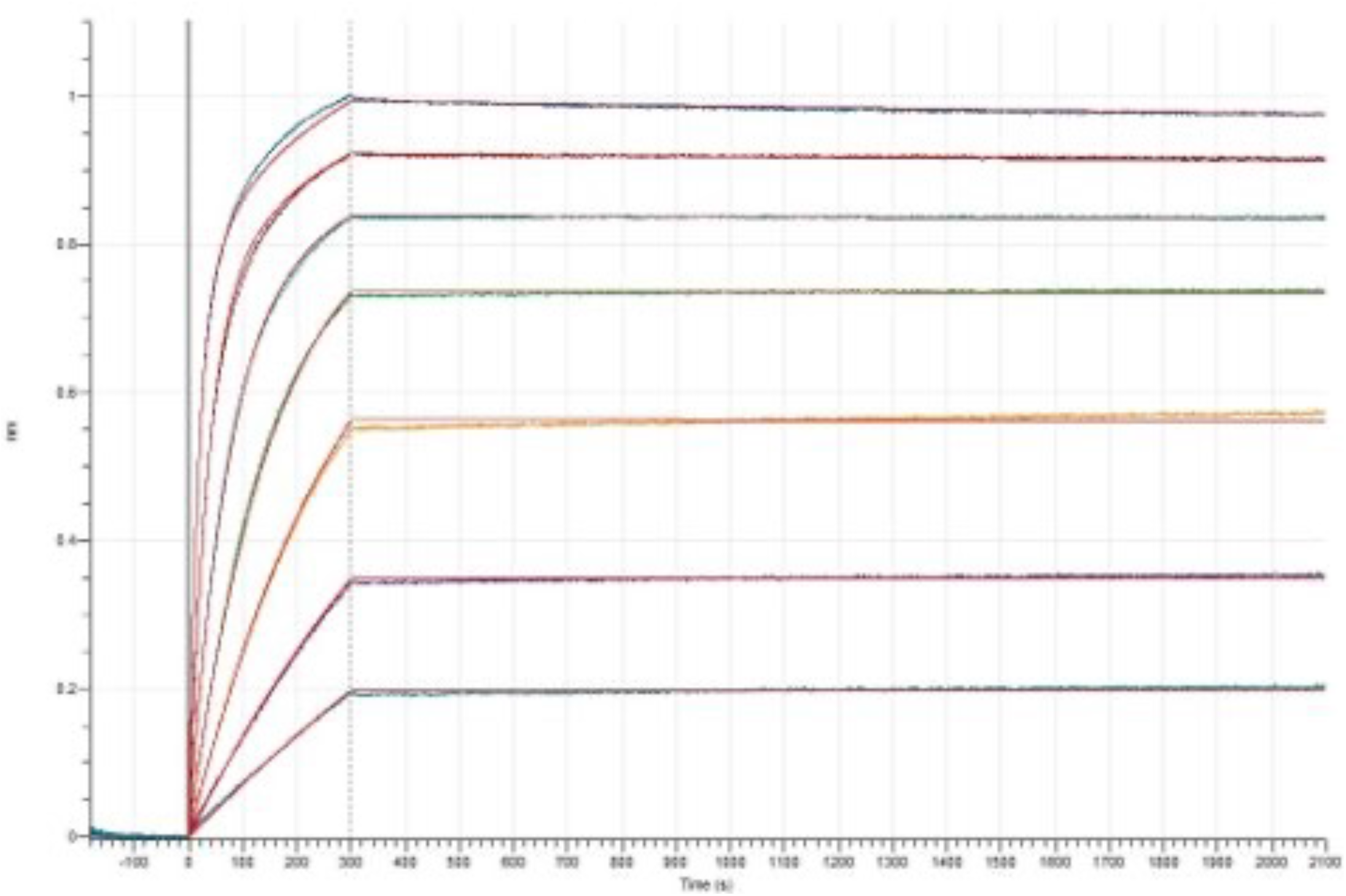
Representative B5-OspC BLI sensorgram. Biotinylated OspC_A_ (3 μg/mL) in buffer (PBS containing 2% w/v BSA) was captured onto streptavidin biosensors, then exposed to a 2-fold serial dilutions (100 to 1.56 nM) for 5 min then 30 min dissociation, as detailed in the Materials and Methods. Results were analyzed using Data Analysis HT 12.0 software, and fit to a 1:2 bivalent analyte model.

**Figure S3.**
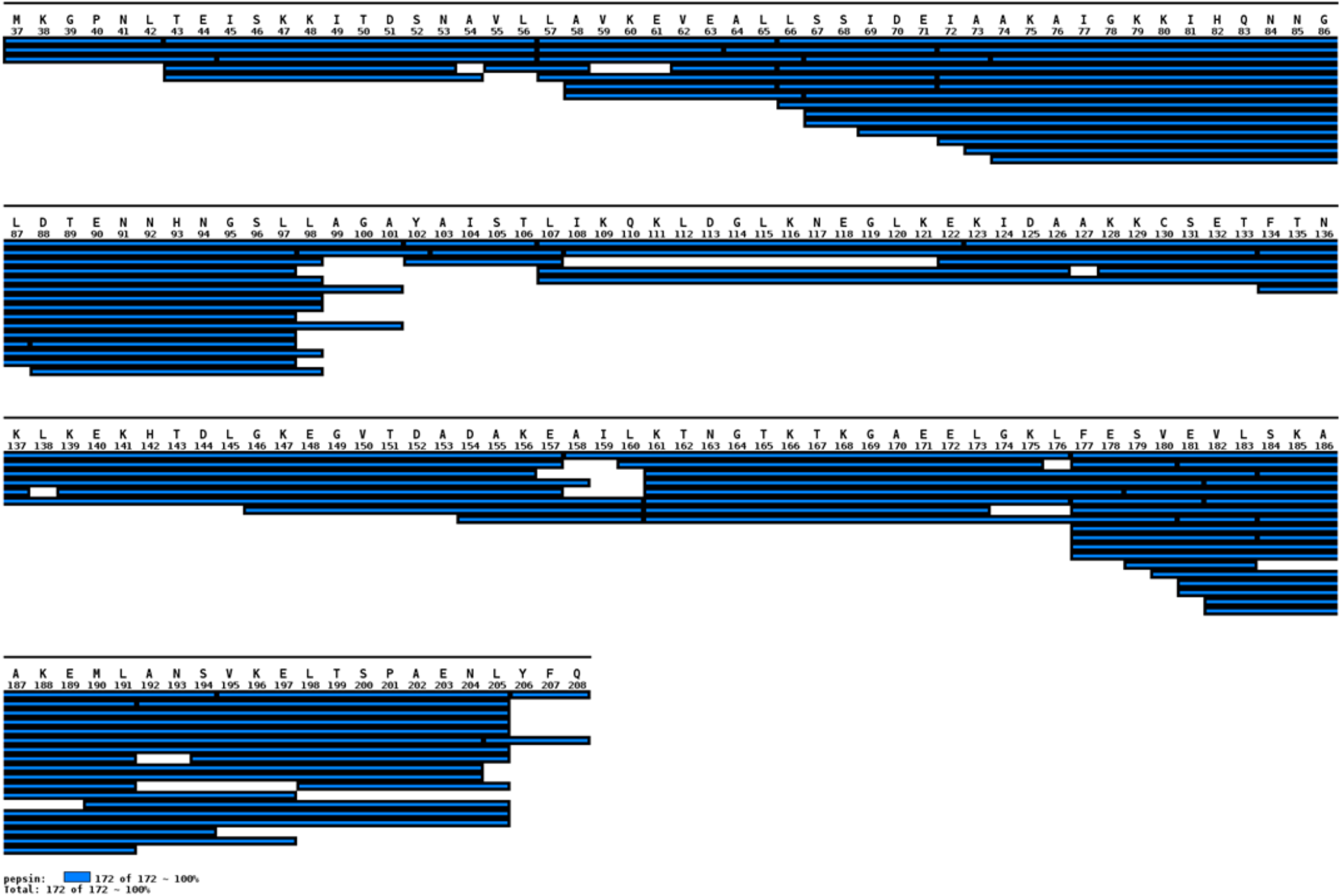
Platformed peptidic map of OspC_A_. OspC_A_ was digested by in house prepared immobilized pepsin column (2.1 ×50 mm). Digested peptides were by trapped and desalted by C-8 column for 120 s and separated by a C-18 column,. For LC, mobile phase A was 0.1% formic acid in water, and B was 0.1% formic acid in acetonitrile. A total of 25 min LC method, 10 minutes with 15%-35%B was used to separate peptides, as described in the Materials and Methods. A total of 87 peptides were identified and mapped for OspC_A_.

**Figure S4.**
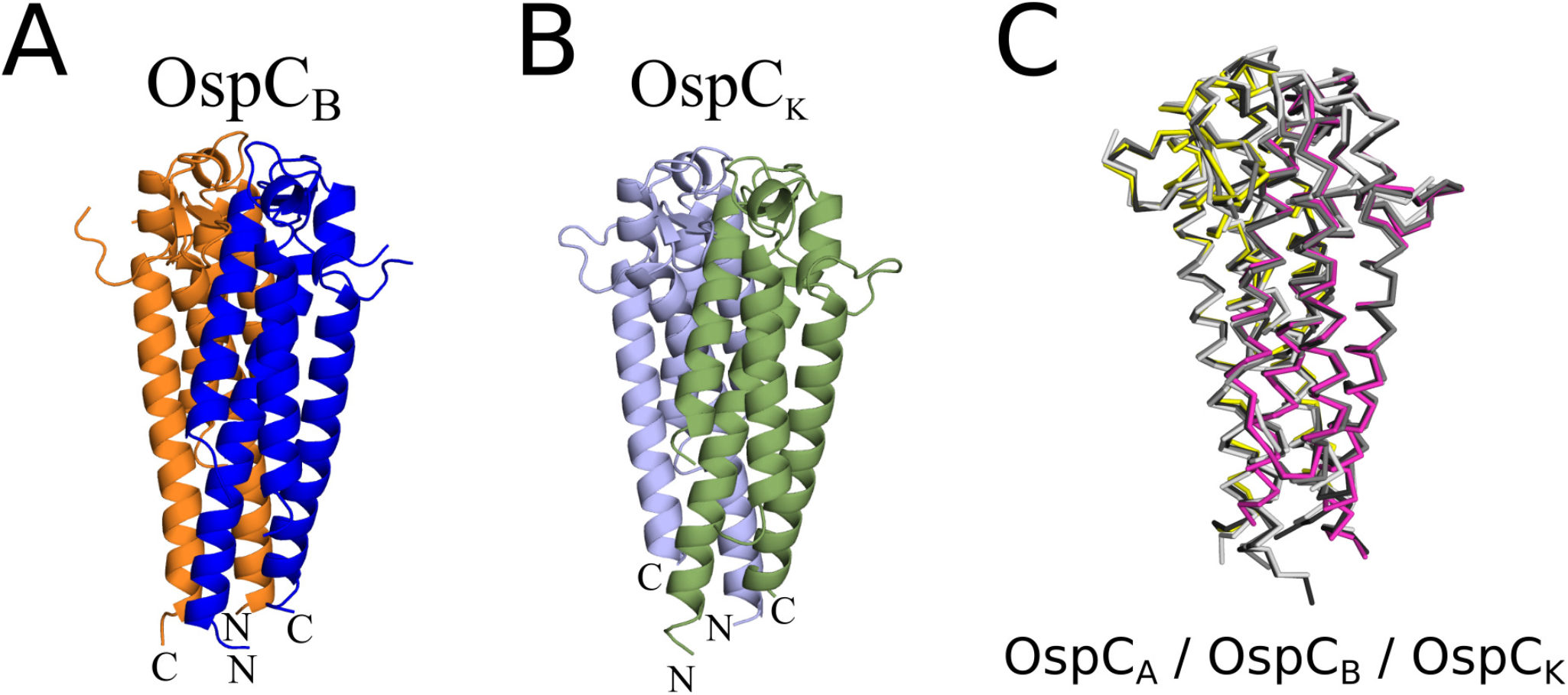
Structural comparison of OspC_B_ and OspC_K_ with OspC_A_. Structures of (A) OspC_B_ (blue and orange), and (B) OspC_K_ (green and slate blue) depicted as ribbon diagrams with N- and C-termini labelled. (C) C-a traces of OspC_A_ homodimer from the B5-OspC_A_ complex (magenta and yellow) superpositioned with unbound OspC_A_ (PDB ID: 1GGQ) colored dark gray, OspCB colored medium gray, and OspCK light gray.

**Figure S5.**
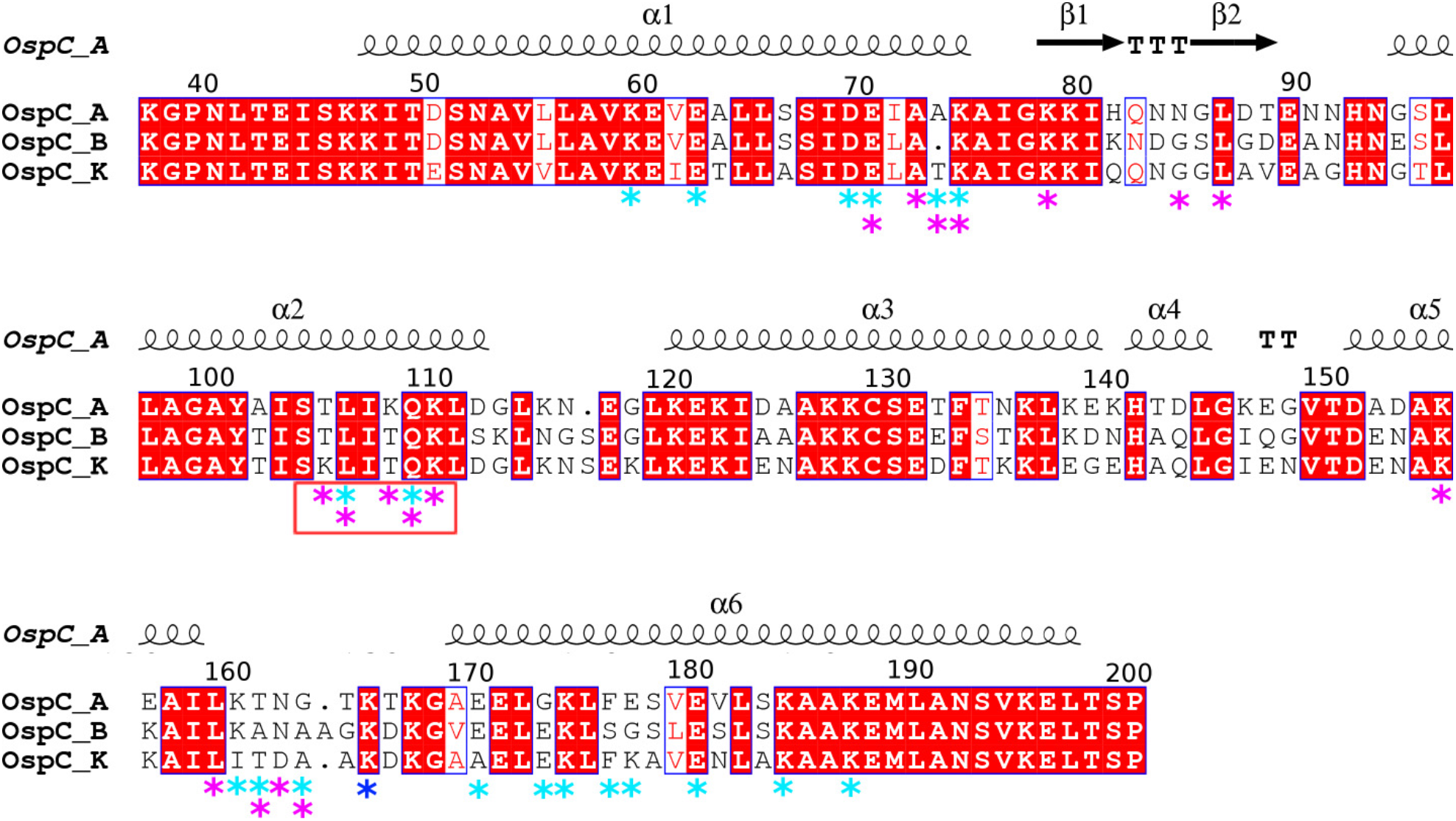
Sequence alignment of OspC A, B, K. Sequence alignment of OspC types A, B, and K from *B. burgdorferi* depicting the B5-interacting region of OspC_A_. Magenta asterisks depict B5 V_L_ (L1-L3) interacting residues, cyan asterisks depict B5 V_H_ (H1-H3) interacting residues, and blue asterisks denote interaction with B5 V_L_ and V_H_ framework region residues. Secondary structural elements from OspC_A_ bound to B5 are illustrated with a-helices 1-6 drawn as black coils and β-strands 1-2 drawn as black arrows above the sequence and labelled accordingly. Red background with white letters connotes sequence identity, white background with red letters connotes sequence similarity. The orange box below sequence region 100-110 shows B5 contact with the second OspC_A_ monomer. Figure made with ClustalW and ESPript 3.0, as described in the Materials and Methods.

**Figure S6.**
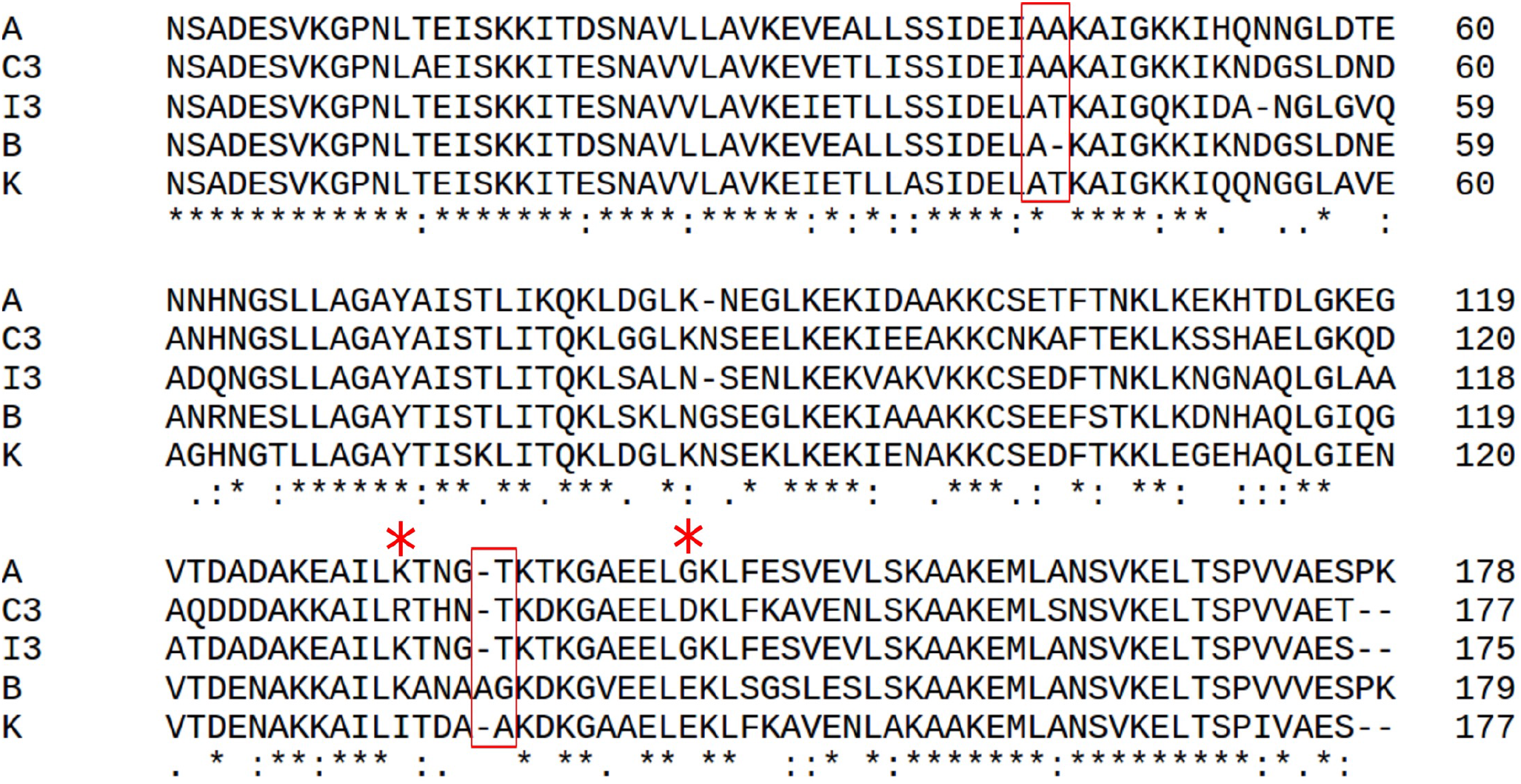
Sequence alignment of OspC types that conceivably bind B5. Primary sequence alignment of OspC types C3 and I3 with OspC_A_, OspC_B_, and OspC_K_ highlighting the key sequence similarities of OspC_C3_ and OspC_I3_ to OspC_A_. Red rectangles encapsulate critical residues in α-helix 1 and loop 5-6 highlighting the insertion at residue 74 and deletion at residue 165 in OspCB that antagonize B5 binding. Red asterisks above the sequence identify the critical OspC_A_ residues 161 and 175 which support interaction with B5. Black asterisks below the sequence denote sequence identity with two dots and one dot showing relatively reduced sequence similarity. Figure made with Clustal Omega.

## References Cited

Adams, P.D., Afonine, P.V., Bunkoczi, G., Chen, V.B., Davis, I.W., Echols, N., Headd, J.J., Hung, L.W., Kapral, G.J., Grosse-Kunstleve, R.W., et al. (2010). PHENIX: a comprehensive Python-based system for macromolecular structure solution. Acta Crystallogr D Biol Crystallogr 66, 213–221. 10.1107/S0907444909052925S0907444909052925 [pii].

Angalakurthi, S.K., Vance, D.J., Rong, Y., Nguyen, C.M.T., Rudolph, M.J., Volkin, D., Middaugh, C.R., Weis, D.D., and Mantis, N.J. (2018). A Collection of Single-Domain Antibodies that Crowd Ricin Toxin’s Active Site. Antibodies (Basel) 7. 10.3390/antib7040045.

Arnaboldi, P.M., Seedarnee, R., Sambir, M., Callister, S.M., Imparato, J.A., and Dattwyler, R.J. (2013). Outer surface protein C peptide derived from Borrelia burgdorferi sensu stricto as a target for serodiagnosis of early lyme disease. Clin Vaccine Immunol 20, 474–481. 10.1128/CVI.00608-12.

Barbour, A.G., and Travinsky, B. (2010). Evolution and distribution of the ospC Gene, a transferable serotype determinant of Borrelia burgdorferi. MBio 1. 10.1128/mBio.00153-10.

Baum, E., Randall, A.Z., Zeller, M., and Barbour, A.G. (2013). Inferring epitopes of a polymorphic antigen amidst broadly cross-reactive antibodies using protein microarrays: a study of OspC proteins of Borrelia burgdorferi. PLoS One 8, e67445. 10.1371/journal.pone.0067445.

Bhatia, B., Hillman, C., Carracoi, V., Cheff, B.N., Tilly, K., and Rosa, P.A. (2018). Infection history of the blood-meal host dictates pathogenic potential of the Lyme disease spirochete within the feeding tick vector. PLoS Pathog 14, e1006959. 10.1371/journal.ppat.1006959.

Bockenstedt, L.K., Hodzic, E., Feng, S., Bourrel, K.W., de Silva, A., Montgomery, R.R., Fikrig, E., Radolf, J.D., and Barthold, S.W. (1997). Borrelia burgdorferi strain-specific Osp C-mediated immunity in mice. Infect Immun 65, 4661–4667. 10.1128/iai.65.11.4661-4667.1997.

Brier, S., Rasetti-Escargueil, C., Wijkhuisen, A., Simon, S., Marechal, M., Lemichez, E., and Popoff, M.R. (2021). Characterization of a highly neutralizing single monoclonal antibody to botulinum neurotoxin type A. FASEB J 35, e21540. 10.1096/fj.202002492R.

Buckles, E.L., Earnhart, C.G., and Marconi, R.T. (2006). Analysis of antibody response in humans to the type A OspC loop 5 domain and assessment of the potential utility of the loop 5 epitope in Lyme disease vaccine development. Clin Vaccine Immunol 13, 1162–1165. 10.1128/CVI.00099-06.

Bunikis, J., Garpmo, U., Tsao, J., Berglund, J., Fish, D., and Barbour, A.G. (2004). Sequence typing reveals extensive strain diversity of the Lyme borreliosis agents Borrelia burgdorferi in North America and Borrelia afzelii in Europe. Microbiology (Reading) 150, 1741–1755. 10.1099/mic.0.26944-0.

Caimano, M.J., Groshong, A.M., Belperron, A., Mao, J., Hawley, K.L., Luthra, A., Graham, D.E., Earnhart, C.G., Marconi, R.T., Bockenstedt, L.K., et al. (2019). The RpoS Gatekeeper in Borrelia burgdorferi: An Invariant Regulatory Scheme That Promotes Spirochete Persistence in Reservoir Hosts and Niche Diversity. Front Microbiol 10, 1923. 10.3389/fmicb.2019.01923.

Caine, J.A., and Coburn, J. (2016). Multifunctional and Redundant Roles of Borrelia burgdorferi Outer Surface Proteins in Tissue Adhesion, Colonization, and Complement Evasion. Front Immunol 7, 442. 10.3389/fimmu.2016.00442.

Chen, G., Karauzum, H., Long, H., Carranza, D., Holtsberg, F.W., Howell, K.A., Abaandou, L., Zhang, B., Jarvik, N., Ye, W., et al. (2019). Potent Neutralization of Staphylococcal Enterotoxin B In Vivo by Antibodies that Block Binding to the T-Cell Receptor. J Mol Biol 431, 4354–4367. 10.1016/j.jmb.2019.03.017.

Dattwyler, R.J., and Gomes-Solecki, M. (2022). The year that shaped the outcome of the OspA vaccine for human Lyme disease. NPJ Vaccines 7, 10. 10.1038/s41541-022-00429-5.

De Silva, A.M., and Fikrig, E. (1995). Growth and migration of Borrelia burgdorferi in Ixodes ticks during blood feeding. Am J Trop Med Hyg 53, 397–404. 10.4269/ajtmh.1995.53.397.

Di, L., Akther, S., Bezrucenkovas, E., Ivanova, L., Sulkow, B., Wu, B., Mneimneh, S., Gomes-Solecki, M., and Qiu, W.G. (2022). Maximum antigen diversification in a lyme bacterial population and evolutionary strategies to overcome pathogen diversity. ISME J 16, 447–464. 10.1038/s41396-021-01089-4.

Earnhart, C.G., Buckles, E.L., Dumler, J.S., and Marconi, R.T. (2005). Demonstration of OspC type diversity in invasive human lyme disease isolates and identification of previously uncharacterized epitopes that define the specificity of the OspC murine antibody response. Infect Immun 73, 7869–7877. 10.1128/IAI.73.12.7869-7877.2005.

Earnhart, C.G., Leblanc, D.V., Alix, K.E., Desrosiers, D.C., Radolf, J.D., and Marconi, R.T. (2010). Identification of residues within ligand-binding domain 1 (LBD1) of the Borrelia burgdorferi OspC protein required for function in the mammalian environment. Mol Microbiol 76, 393–408. 10.1111/j.1365-2958.2010.07103.x.

Earnhart, C.G., and Marconi, R.T. (2007). OspC phylogenetic analyses support the feasibility of a broadly protective polyvalent chimeric Lyme disease vaccine. Clin Vaccine Immunol 14, 628–634. 10.1128/CVI.00409-06.

Edmondson, D.G., Prabhakaran, S., Norris, S.J., Ullmann, A.J., Piesman, J., Dolan, M., Probst, C., Radzimski, C., Stöcker, W., and Komorowski, L. (2017). Enhanced Protective Immunogenicity of Homodimeric Borrelia burgdorferi Outer Surface Protein C. Clin Vaccine Immunol 24. 10.1128/cvi.00306-16.

Eicken, C., Sharma, V., Klabunde, T., Lawrenz, M.B., Hardham, J.M., Norris, S.J., and Sacchettini, J.C. (2002). Crystal structure of Lyme disease variable surface antigen VlsE of Borrelia burgdorferi. J Biol Chem 277, 21691–21696. 10.1074/jbc.M201547200.

Eicken, C., Sharma, V., Klabunde, T., Owens, R.T., Pikas, D.S., Hook, M., and Sacchettini, J.C. (2001). Crystal structure of Lyme disease antigen outer surface protein C from Borrelia burgdorferi. J Biol Chem 276, 10010–10015. 10.1074/jbc.M010062200.

Emsley, P., Lohkamp, B., Scott, W.G., and Cowtan, K. (2010). Features and development of Coot. Acta Crystallogr D Biol Crystallogr 66, 486–501. 10.1107/S0907444910007493S0907444910007493 [pii].

Gilmore, R.D., Jr., Kappel, K.J., Dolan, M.C., Burkot, T.R., and Johnson, B.J. (1996). Outer surface protein C (OspC), but not P39, is a protective immunogen against a tick-transmitted Borrelia burgdorferi challenge: evidence for a conformational protective epitope in OspC. Infect Immun 64, 2234–2239. 10.1128/iai.64.6.2234-2239.1996.

Gilmore, R.D., Jr., and Mbow, M.L. (1999). Conformational nature of the Borrelia burgdorferi B31 outer surface protein C protective epitope. Infect Immun 67, 5463–5469. 10.1128/IAI.67.10.5463-5469.1999.

Gilmore, R.D., Jr., and Piesman, J. (2000). Inhibition of Borrelia burgdorferi migration from the midgut to the salivary glands following feeding by ticks on OspC-immunized mice. Infect Immun 68, 411–414. 10.1128/IAI.68.1.411-414.2000.

Gomes-Solecki, M., Arnaboldi, P.M., Backenson, P.B., Benach, J.L., Cooper, C.L., Dattwyler, R.J., Diuk-Wasser, M., Fikrig, E., Hovius, J.W., Laegreid, W., et al. (2020). Protective Immunity and New Vaccines for Lyme Disease. Clin Infect Dis 70, 1768–1773. 10.1093/cid/ciz872.

Grauslund, L.R., Calvaresi, V., Pansegrau, W., Norais, N., and Rand, K.D. (2021). Epitope and Paratope Mapping by HDX-MS Combined with SPR Elucidates the Difference in Bactericidal Activity of Two Anti-NadA Monoclonal Antibodies. J Am Soc Mass Spectrom 32, 1575–1582. 10.1021/jasms.0c00431.

Hanincova, K., Mukherjee, P., Ogden, N.H., Margos, G., Wormser, G.P., Reed, K.D., Meece, J.K., Vandermause, M.F., and Schwartz, I. (2013). Multilocus sequence typing of Borrelia burgdorferi suggests existence of lineages with differential pathogenic properties in humans. PLoS One 8, e73066. 10.1371/journal.pone.0073066.

Haque, H.M.E., Ejemel, M., Vance, D.J., Willsey, G., Rudolph, M.J., Cavacini, L.A., Wang, Y., Mantis, N.J., and Weis, D.D. (2022). Human B Cell Epitope Map of the Lyme Disease Vaccine Antigen, OspA. ACS Infect Dis. 10.1021/acsinfecdis.2c00346.

Hodge, E.A., Naika, G.S., Kephart, S.M., Nguyen, A., Zhu, R., Benhaim, M.A., Guo, W., Moore, J.P., Hu, S.L., Sanders, R.W., and Lee, K.K. (2022). Structural dynamics reveal isolate-specific differences at neutralization epitopes on HIV Env. iScience 25, 104449. 10.1016/j.isci.2022.104449.

Izac, J.R., Camire, A.C., Earnhart, C.G., Embers, M.E., Funk, R.A., Breitschwerdt, E.B., and Marconi, R.T. (2019). Analysis of the antigenic determinants of the OspC protein of the Lyme disease spirochetes: Evidence that the C10 motif is not immunodominant or required to elicit bactericidal antibody responses. Vaccine 37, 2401–2407. 10.1016/j.vaccine.2019.02.007.

Izac, J.R., and Marconi, R.T. (2019). Diversity of the Lyme Disease Spirochetes and its Influence on Immune Responses to Infection and Vaccination. Vet Clin North Am Small Anim Pract 49, 671–686. 10.1016/j.cvsm.2019.02.007.

Izac, J.R., O’Bier, N.S., Oliver, L.D., Jr., Camire, A.C., Earnhart, C.G., LeBlanc Rhodes, D.V., Young, B.F., Parnham, S.R., Davies, C., and Marconi, R.T. (2020). Development and optimization of OspC chimeritope vaccinogens for Lyme disease. Vaccine 38, 1915–1924. 10.1016/j.vaccine.2020.01.027.

Kugeler, K.J., Schwartz, A.M., Delorey, M.J., Mead, P.S., and Hinckley, A.F. (2021). Estimating the Frequency of Lyme Disease Diagnoses, United States, 2010-2018. Emerg Infect Dis 27, 616–619. 10.3201/eid2702.202731.

Kumaran, D., Eswaramoorthy, S., Luft, B.J., Koide, S., Dunn, J.J., Lawson, C.L., and Swaminathan, S. (2001). Crystal structure of outer surface protein C (OspC) from the Lyme disease spirochete, Borrelia burgdorferi. EMBO J 20, 971–978. 10.1093/emboj/20.5.971.

Malito, E., Faleri, A., Lo Surdo, P., Veggi, D., Maruggi, G., Grassi, E., Cartocci, E., Bertoldi, I., Genovese, A., Santini, L., et al. (2013). Defining a protective epitope on factor H binding protein, a key meningococcal virulence factor and vaccine antigen. Proc Natl Acad Sci U S A 110, 3304–3309. 10.1073/pnas.12228451101222845110 [pii].

Mathiesen, M.J., Holm, A., Christiansen, M., Blom, J., Hansen, K., Ostergaard, S., and Theisen, M. (1998). The dominant epitope of Borrelia garinii outer surface protein C recognized by sera from patients with neuroborreliosis has a surface-exposed conserved structural motif. Infect Immun 66, 4073–4079. 10.1128/IAI.66.9.4073-4079.1998.

Mbow, M.L., Gilmore, R.D., Jr., Stevenson, B., Golde, W.T., Piesman, J., and Johnson, B.J. (2002). Borrelia burgdorferi-specific monoclonal antibodies derived from mice primed with Lyme disease spirochete-infected Ixodes scapularis ticks. Hybrid Hybridomics 21, 179–182. 10.1089/153685902760173890.

Mbow, M.L., Gilmore, R.D., Jr., and Titus, R.G. (1999). An OspC-specific monoclonal antibody passively protects mice from tick-transmitted infection by Borrelia burgdorferi B31. Infect Immun 67, 5470–5472. 10.1128/IAI.67.10.5470-5472.1999.

Nadelman, R.B., Hanincova, K., Mukherjee, P., Liveris, D., Nowakowski, J., McKenna, D., Brisson, D., Cooper, D., Bittker, S., Madison, G., et al. (2012). Differentiation of reinfection from relapse in recurrent Lyme disease. N Engl J Med 367, 1883–1890. 10.1056/NEJMoa1114362.

Norek, A., and Janda, L. (2017). Epitope mapping of Borrelia burgdorferi OspC protein in homodimeric fold. Protein Sci 26, 796–806. 10.1002/pro.3125.

Ohnishi, J., Piesman, J., and de Silva, A.M. (2001). Antigenic and genetic heterogeneity of Borrelia burgdorferi populations transmitted by ticks. Proc Natl Acad Sci U S A 98, 670–675. 10.1073/pnas.98.2.670.

Otwinowski, Z., and Minor, W. (1997). Processing of x-ray diffraction data collected in oscillation mode.. Methods in Enzymology 276, 307–326.

Pal, U., Yang, X., Chen, M., Bockenstedt, L.K., Anderson, J.F., Flavell, R.A., Norgard, M.V., and Fikrig, E. (2004). OspC facilitates Borrelia burgdorferi invasion of Ixodes scapularis salivary glands. J Clin Invest 113, 220–230. 10.1172/JCI19894.

Preac-Mursic, V., Wilske, B., Patsouris, E., Jauris, S., Will, G., Soutschek, E., Rainhardt, S., Lehnert, G., Klockmann, U., and Mehraein, P. (1992). Active immunization with pC protein of Borrelia burgdorferi protects gerbils against B. burgdorferi infection. Infection 20, 342–349. 10.1007/BF01710681.

Probert, W.S., and LeFebvre, R.B. (1994). Protection of C3H/HeN mice from challenge with Borrelia burgdorferi through active immunization with OspA, OspB, or OspC, but not with OspD or the 83-kilodalton antigen. Infect Immun 62, 1920–1926. 10.1128/iai.62.5.1920-1926.1994.

Pulzova, L., Flachbartova, Z., Bencurova, E., Potocnakova, L., Comor, L., Schreterova, E., and Bhide, M. (2016). Identification of B-cell epitopes of Borrelia burgdorferi outer surface protein C by screening a phage-displayed gene fragment library. Microbiol Immunol 60, 669–677. 10.1111/1348-0421.12438.

Radolf, J.D., Strle, K., Lemieux, J.E., and Strle, F. (2021). Lyme Disease in Humans. Curr Issues Mol Biol 42, 333–384. 10.21775/cimb.042.333.

Rana, V.S., Kitsou, C., Dumler, J.S., and Pal, U. (2022). Immune evasion strategies of major tick-transmitted bacterial pathogens. Trends Microbiol. 10.1016/j.tim.2022.08.002.

Rong, Y., Torres-Velez, F.J., Ehrbar, D., Doering, J., Song, R., and Mantis, N.J. (2020). An intranasally administered monoclonal antibody cocktail abrogates ricin toxin-induced pulmonary tissue damage and inflammation. Hum Vaccin Immunother 16, 793–807. 10.1080/21645515.2019.1664243.

Rousselle, J.C., Callister, S.M., Schell, R.F., Lovrich, S.D., Jobe, D.A., Marks, J.A., and Wieneke, C.A. (1998). Borreliacidal antibody production against outer surface protein C of Borrelia burgdorferi. J Infect Dis 178, 733–741.

Schwan, T.G., Piesman, J., Golde, W.T., Dolan, M.C., and Rosa, P.A. (1995). Induction of an outer surface protein on Borrelia burgdorferi during tick feeding. Proc Natl Acad Sci U S A 92, 2909–2913. 10.1073/pnas.92.7.2909.

Seinost, G., Dykhuizen, D.E., Dattwyler, R.J., Golde, W.T., Dunn, J.J., Wang, I.N., Wormser, G.P., Schriefer, M.E., and Luft, B.J. (1999). Four clones of Borrelia burgdorferi sensu stricto cause invasive infection in humans. Infect Immun 67, 3518–3524.

Seow, J., Khan, H., Rosa, A., Calvaresi, V., Graham, C., Pickering, S., Pye, V.E., Cronin, N.B., Huettner, I., Malim, M.H., et al. (2022). A neutralizing epitope on the SD1 domain of SARS-CoV-2 spike targeted following infection and vaccination. Cell Rep 40, 111276. 10.1016/j.celrep.2022.111276.

Steere, A.C., Strle, F., Wormser, G.P., Hu, L.T., Branda, J.A., Hovius, J.W., Li, X., and Mead, P.S. (2016). Lyme borreliosis. Nat Rev Dis Primers 2, 16090. 10.1038/nrdp.2016.90.

Vinciauskaite, V., and Masson, G.R. (2022). Fundamentals of HDX-MS. Essays Biochem. 10.1042/EBC20220111.

Wang, I.N., Dykhuizen, D.E., Qiu, W., Dunn, J.J., Bosler, E.M., and Luft, B.J. (1999). Genetic diversity of ospC in a local population of Borrelia burgdorferi sensu stricto. Genetics 151, 15–30. 10.1093/genetics/151.1.15.

Wang, L., Pan, H., and Smith, D.L. (2002). Hydrogen exchange-mass spectrometry: optimization of digestion conditions. Mol Cell Proteomics 1, 132–138.

Wang, Y., Kern, A., Boatright, N.K., Schiller, Z.A., Sadowski, A., Ejemel, M., Souders, C.A., Reimann, K.A., Hu, L., Thomas, W.D., Jr., and Klempner, M.S. (2016). Pre-exposure Prophylaxis With OspA-Specific Human Monoclonal Antibodies Protects Mice Against Tick Transmission of Lyme Disease Spirochetes. J Infect Dis 214, 205–211. 10.1093/infdis/jiw151.

Ward, A.B., and Wilson, I.A. (2020). Innovations in structure-based antigen design and immune monitoring for next generation vaccines. Curr Opin Immunol 65, 50–56. 10.1016/j.coi.2020.03.013.

Wilske, B., Luft, B., Schubach, W.H., Zumstein, G., Jauris, S., Preac-Mursic, V., and Kramer, M.D. (1992). Molecular analysis of the outer surface protein A (OspA) of Borrelia burgdorferi for conserved and variable antibody binding domains. Med Microbiol Immunol 181, 191–207. 10.1007/bf00215765.

Winn, M.D., Ballard, C.C., Cowtan, K.D., Dodson, E.J., Emsley, P., Evans, P.R., Keegan, R.M., Krissinel, E.B., Leslie, A.G., McCoy, A., et al. (2011). Overview of the CCP4 suite and current developments. Acta Crystallogr D Biol Crystallogr 67, 235–242. 10.1107/S0907444910045749S0907444910045749 [pii].

Wormser, G.P. (2022). A brief history of OspA vaccines including their impact on diagnostic testing for Lyme disease. Diagn Microbiol Infect Dis 102, 115572. 10.1016/j.diagmicrobio.2021.115572.

Yu, Z., Carter, J.M., Sigal, L.H., and Stein, S. (1996). Multi-well ELISA based on independent peptide antigens for antibody capture. Application to Lyme disease serodiagnosis. J Immunol Methods 198, 25–33.

Zhong, W., Stehle, T., Museteanu, C., Siebers, A., Gern, L., Kramer, M., Wallich, R., and Simon, M.M. (1997). Therapeutic passive vaccination against chronic Lyme disease in mice. Proc Natl Acad Sci U S A 94, 12533–12538. 10.1073/pnas.94.23.12533.

Zuckert, W.R., Kerentseva, T.A., Lawson, C.L., and Barbour, A.G. (2001). Structural conservation of neurotropism-associated VspA within the variable Borrelia Vsp-OspC lipoprotein family. J Biol Chem 276, 457–463. 10.1074/jbc.M008449200.

